# Environmental DNA reveals biotic homogenization alongside elevated native and non-native species richness in marinas, supporting an ecological portuarization syndrome

**DOI:** 10.1101/2025.10.27.684730

**Authors:** Ginevra Lilli, Annaëlle Caillarec-Joly, Clément Violet, Marc Bouchoucha, Xavier Turon, Sophie Arnaud-Haond, Frédérique Viard

## Abstract

The global expansion of ports has created novel marine urban ecosystems that remain understudied. Recent evidence suggests that ports promote distinct evolutionary trajectories due to their unique environmental conditions and the continuous introduction of non-indigenous species (NIS), a phenomenon previously coined the “biological portuarization syndrome”. In this study, we examine the ecological dimension of this concept by using environmental DNA (eDNA) metabarcoding, to compare animal assemblages in 12 recreational ports (i.e., marinas) with those found in adjacent natural habitats in the north-western Mediterranean Sea. By combining three markers, we identified 536 species belonging to 18 phyla, including 50 species classified as non-indigenous or cryptogenic based on previous records from traditional surveys. Our results reveal marked differences in both taxonomic and functional composition between marinas and natural communities. Notably, contrary to expectations of reduced species richness due to environmental filtering in marinas, we observed higher species richness in these habitats. This was true even when considering native species alone, and it was broadly consistent across phyla: nine phyla exhibited significantly higher species richness in marinas, eight showed no significant difference between habitats, and only one (Arthropoda) had higher species richness in natural habitats. Furthermore, consistent with the hypothesis of biotic homogenization in anthropogenic habitats, animal assemblages were more homogeneous among marinas than among natural habitats. Overall, our findings show the impact of human-made habitats on regional biodiversity patterns, and suggest that the biological portuarization syndrome extends beyond evolutionary processes to include distinctive ecological patterns.

## Introduction

Since the beginning of the nineteenth century, urban areas and megacities have increased in size and number as part of the urban transition (Day et al., 2021), notably affecting coastal zones (Creel, 2003). Despite their defined borders, urban areas significantly impact the surrounding environment (Elmqvist et al., 2013) and create modified or entirely new habitats. The ecological and evolutionary processes of these habitats are only just beginning to be understood through the field of “urban science” (McKinney, 2002), which has been so far largely focused on terrestrial ecosystems. However, coastal urbanization substantially impacts marine coastal habitats and ecosystems (Bishop et al., 2017; Mayer-Pinto et al., 2018), which calls for strengthening marine urban sciences (Todd et al., 2019). In addition to their key ecological roles, coastal ecosystems support numerous economic sectors, including fisheries, tourism, wind energy production, and mineral resource extraction (Martínez et al., 2007; Seitz et al., 2014; Sempere-Valverde et al., 2022), leading to the worldwide proliferation of ports (Jouffray et al., 2020).

Ports, whether large commercial harbours or small recreational marinas, provide new marine habitats characterized by particular abiotic properties (Alter et al., 2021; Touchard et al., 2023). For example, they are characterized by a high concentration of waste and pollutants, turbidity and an irregular light regime (Cassi et al., 2008; Cutroneo et al., 2017; Schaefer et al., 2025). The same applies to their biotic properties, which remain far less well understood than their abiotic characteristics (Madon et al., 2023). Ports are nevertheless recognized as entry points for non-indigenous species (NIS), which account for a large proportion of the species present in ports (e.g., Ferrario et al., 2017; O’Shaughnessy et al., 2020). The escape of NIS from ports is recognised as one of the main drivers of their spread at regional level into the natural environment (Afonso et al., 2020, Outinen et al. 2021; Ashton et al., 2022). However, NIS occurrence in natural habitats surrounding ports has only been marginally investigated (Soares et al., 2018; Sghaier et al., 2019; Diem et al., 2023; Zarcero et al., 2026), as compared to their occurrence in ports (Tempesti et al., 2020). NIS monitoring in ports indeed supports compliance with regulations, such as the European Marine Strategy Framework Directive and of biosecurity measures, as recommended by the International Maritime Organization guidelines against NIS spread by biofouling (Outinen et al., 2021).

Besides NIS, the overall species richness in ports is another biotic feature requiring special attention (Tanasovici et al., 2025; Pavard et al. 2026). The prevailing view is that species richness is generally lower in these human-made ecosystems than in nearby natural habitats, a pattern commonly attributed to the low tolerance of many species to anthropogenic disturbances (Chapman, 2003; Bulleri & Chapman 2010, Wakefield et al 2013, Schaefer et al., 2025; Zarcero et al., 2026). However, growing evidence suggests that this pattern is not yet generalizable across taxa or environmental settings. For example, fish biodiversity can be similar, or even higher, in marinas than in nearby natural habitats as marinas may provide opportunities for fish foraging, escape from predators, and nursery grounds (Bouchoucha et al., 2016; Manel et al., 2024; Macé et al., 2024, 2026). Similarly, Pavard et al. (2026) observed high species richness of foraminiferal communities and macrofaunal assemblages in sediments form the heavily polluted port of Dunkirk. Although no reference sites outside the port were included, the authors noted that species richness was higher than reported in studies of marinas and natural sites in the region. Together, these findings suggest that the relationship between port environments and species richness may be more context-dependent than generally assumed and warrants further investigation.

Port communities have also been shown to be characterized by stress-tolerant, synanthropic, and non-indigenous species, which are “filtered” by port abiotic properties (Tempesti et al., 2022; Pavard et al., 2026). Therefore, besides structural and functional distinctiveness as compared to nearby natural habitats, we expect communities within ports to resemble each other more than communities outside of ports – i.e., biotic homogenization (Olden et al., 2004; Todd et al., 2019). These global consequences have only been marginally explored (McKinney & Lockwood, 1999; Lokatis & Jeschke, 2022).

The singularity of ports, acting as replicated highly-modified environments, led Touchard et al. (2023) to define a “biological portuarization” syndrome as “*the repeated evolution of marine species in port ecosystems under human-altered selective pressures*”. The present work aims to further investigate the portuarization syndrome from an ecological perspective and explore the aforementioned gaps on animal marine communities, by comparing recreational ports (i.e. marinas) and their nearby natural habitats at a regional scale, in the northwestern Mediterranean Sea. We selected marinas because, while sharing the distinct characteristic of ports described above, their abundance, properties and accessibility enabled us to study replicated situations in the study region. Due to the growing recreational boating industry, the number of small ports in the Mediterranean Sea has increased in recent years. Nowadays, there are more than 900 marinas in this basin, over half of which located in the northwest Mediterranean Sea (Carreño & Lloret, 2021). In our study area along the coast of France (continental and insular, including Corsica), 65 recreational ports with 39,900 berths were reported in 2018 (MTES, 2018), providing numerous sailing opportunities and port connections. Mediterranean marinas are indeed characterized by intense maritime traffic connecting them at various scales (Ulman et al., 2019). Additionally, previous studies have shown the high prevalence of NIS in these habitats (Ulman et al., 2017). However, the diversity of native species is comparatively poorly known, leaving a gap in our understanding of the overall biodiversity inhabiting marinas. This knowledge gap partly stems from the traditional visual methods used for investigating marine biodiversity, which often involve a trade-off between taxonomic resolution and breadth. This restricts investigations to either a few selected taxa across multiple sites or to multiple taxa in a limited spatial context. Environmental DNA (eDNA) metabarcoding is increasingly used to overcome these limitations and complement traditional methods for biodiversity assessment as well as NIS detection (Stat et al., 2017; Couton et al. 2022; Keck et al., 2022; Fonseca et al., 2023; Dafforn et al., 2025; Zarcero et al., 2026). Additionally, as our aim was to compare multiple marinas with their adjacent habitats, it was essential to ensure consistency in sampling and data collection across a large number of sites, at a regional level. This is a requirement that is difficult to achieve with traditional survey methods, but can be attained with eDNA metabarcoding. This is a highly standardized approach, provided the entire process, from field sampling to bioinformatic data analysis, is conducted by the same operator (Baird and Hajibabaei 2012; Blackman et al. 2024).

We used eDNA metabarcoding of seawater samples to assess the diversity and composition of animal communities in 12 pairs of northwestern Mediterranean marinas and neighbouring natural areas. Based on the previous studies cited above, we tested the following hypotheses: species assemblages in ports differ from those in natural habitats (H1) and show biotic homogenization across a regional scale (H2); communities in ports are characterized by greater abundance of NIS (H3), lower species richness than natural habitats (H4), and taxonomic and functional traits distinct from communities from the nearby natural habitats (H5).

## Materials and Methods

### Data collection

Seawater samples used for this study were collected during summer 2021 (June-August) within the framework of the SUCHIMed campaign (https://doi.org/10.17600/18001619), in 12 marinas along the French Mediterranean continental coast and five along the Corsican coast (Figure 1A; additional information about the marinas selected for the study are reported in the Appendix S1.1). In contrast to many large commercial ports, or even marinas in other seas under strong tidal regime (e.g., English Channel), all the study marinas are directly connected to the open coast with no gates, locks, or other structures restricting water exchange. Seawater was collected in triplicates from three pontoons in each of the 12 marinas. Before entering the marina, samples were also collected in triplicates from a boat in three spots located within a few hundred meters from the marina entrance. Two litres of seawater were collected at each sampling site over a maximum depth of 50 cm and immediately filtered on site or on boat by using 0.22 µm Sterivex filters and a VAMPIRE peristaltic pump (Bürkle GmbH, Deutschland). Filters were filled with 2 mL of preservation buffer (sucrose 0.75 M, Tris 0.05 M pH 8, EDTA 0.04 M) and stored at -20°C until DNA extraction. In total, 74 samples, including two field controls obtained by filtering ultrapure water with the equipment used on site, were collected across the 12 localities.

**Figure 1:**
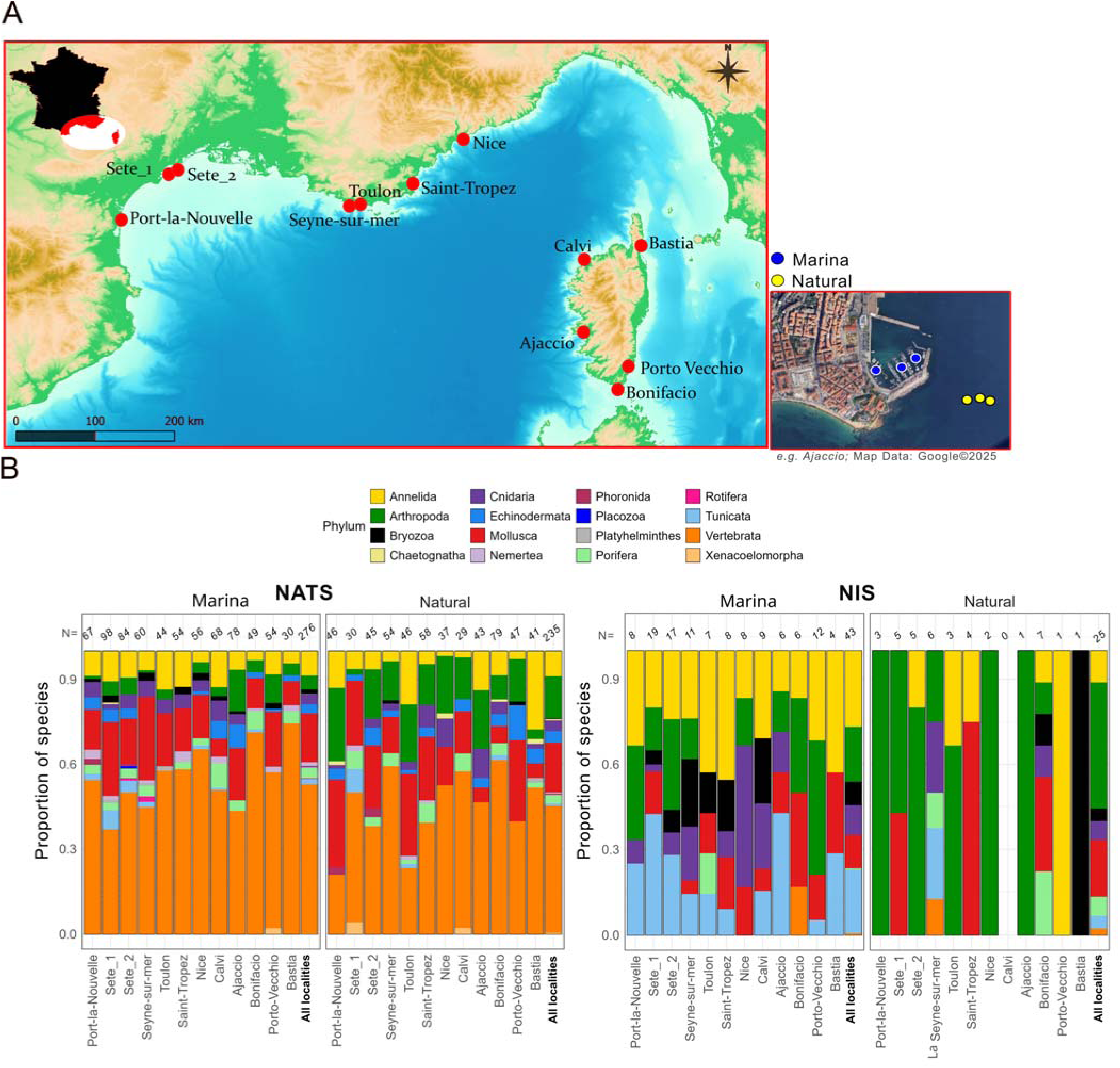
Sampling locations and taxonomic composition of native and non-indigenous species per habitat and locality. A) French localities where seawater was collected at three sites inside marina and three site in nearby natural habitats. An example is shown for Ajaccio in the small panel. Basemap: GEBCO_Grid, colour-shaded for depth (GEBCO Compilation Group, 2023); B) Proportion of native species (NATS, left panel) and NIS (right panel) per phylum per habitat type (i.e. Marina vs Natural habitat) by each locality and across all localities. The number of NATS and NIS recorded in the different habitat types is reported at the top of each bar. The proportions of species included neither in the NATS or NIS categories (“Uncertain” species – see Material & Methods) is reported in Appendix S.1.4 Figure S2.

DNA was extracted from Sterivex filters using the NucleoSnap® Finisher Midi extraction kit (Macherey-Nagel) following the protocol detailed in Couton et al. (2022). One DNA extraction control was included in each extraction series by processing an unused sterivex as those used in the field. PCR amplification was performed on samples and field controls at Station Ifremer (Sète) in a laboratory dedicated to environmental DNA analyses. Each DNA sample was amplified 9 times with each of three primer pairs, selected based on the Animalia target: i) primers mlCOIintF and jgHCO2198 (Leray et al., 2013) to amplify a ca. 313-base-pair (bp) region of subunit I of the cytochrome c oxidase (COI) gene; ii) primers SSU_F04 and SSU_R22mod (Sinniger et al., 2016) to amplify a ca. 450 bp fragment of the V1-V2 region of the 18S nuclear gene and iii) Teleo_04 primers, specifically designed to target teleost fish by amplifying a ca. 160bp region of 12S mitochondrial gene (primer design by Pierre Taberlet; previously used by Roblet et al. (2024)). PCR reactions were carried out in a 10 μL reaction volume. For 18S and 12S, PCR was performed using Q5® High-Fidelity DNA polymerase (New England Biolabs®, Inc.). As the jgHCO2198 forward primer contains inosine, the Qiagen® Multiplex PCR Plus Kit was used for COI instead of a proof-reading polymerase. Negative PCR controls were included in each PCR trials. The first purified PCR products were used for a second PCR with primers including sample tags and indexes.

Library preparation was performed at the GenSeq Core Service (University of Montpellier, France), with one library per marker. The libraries were sequenced (2 x 250bp paired-end) by the Novogene company, using the NovaSeq 6000 SP Reagent Kit (500 cycles) and a Novaseq 6000 SP flow cell sequencer.

### Bioinformatic analyses and taxonomic assignment

Paired-end raw reads were pre-processed using cutadapt v1.16 and bbMap v38.22, followed by a modified Unix shell script based on Brandt et al. (2021) (https://gitlab.ifremer.fr/abyss-project/abyss-pipeline) for denoising (DADA2 v1.10) (Callahan et al., 2016) and clustering (Swarm v3.1.0 with d=1 for 12S and 18S and d=3 for COI) (Mahé et al., 2014). OTU tables were decontaminated using the R package decontam prevalence-based method (method = “prevalence”, that identifies contaminant OTUs by comparing their prevalence in true samples versus controls) (Davis et al., 2018). A correction for abundances was applied as done in Wangensteen et al., (2018), with samples sorted by abundance of each OTU and the reads of the samples corresponding to a cumulative frequency of less than 3% of each OTU removed. For COI data, mumu (Mahé, 2025), adapted and modified from Lulu (Frøslev et al., 2017) was applied (with identity 84%, co-occurrence 0.90 and min ratio 0.01) to minimize the bias due to the existence of paralogs and pseudogenes (particularly nuclear degenerated copies of the mitochondrial gene, *numts*). OTU sequences were aligned with those in reference databases using BLAST v2.12.0 (Altschul et al., 1990). Sequence coverage was set at a minimum of 70% for 12S and 80% for COI and 18S. Minimum identity percentages with sequences in reference databases were set following results of Couton et al., (2022): 92% for COI, 90% for 12S (MIDORI2 database) and 99% for 18S (Silva v138 database). OTU tables were rarefied to 100.000 reads per sample, unless the minimum number of reads was higher (i.e., 107,876 for 18S). Prior to running rarefaction, seven samples from the 12S and COI datasets were excluded from further analyses since they displayed fewer than 100,000 reads. Specifically, these were: one sample from the marinas of Bastia, Nice and Port-La-Nouvelle for the 12S dataset; one sample from the marinas of Sete_1 and Sete_2, and one sample from the natural sites in Sete_2 and in Toulon for the COI dataset. Removing these samples resulted in a total of 65 samples common among the three marker libraries and eventually used for the analyses. WoRMS package version 0.4.3 in R (Chamberlain & Vanhoorne, 2023) was used to correct the nomenclature and remove non-marine taxa; Chordata were split into Vertebrata and Tunicata due to the important ecological differences between these two subphyla.

The OTU tables generated from processing the three marker libraries were converted into presence-absence matrices. In order to assign a status of native vs. non-native species to the identified taxa, the matrices were filtered to retain only those OTUs identified at the species level. Next, to obtain the most comprehensive species-level taxonomic table possible, we took advantage of the complementary nature of the markers in terms of taxonomic assignment by merging the species-level assignments from the three single-marker tables into a single table (i.e., creating a Species occurrence table).

### Classification of species as native or non-indigenous

The species dataset (hereafter “COMM” dataset, i.e., whole community) was screened to identify non-indigenous species (NIS). We adopted stringent criteria to the assignment of the native and non-native species status.

Identified species were included as NIS only if those species had already been listed as introduced in European seas, and we also recorded if they had been already reported specifically in the Mediterranean Sea. This assignment was made using the NIS species list reported in Massé et al., (2023), the most recent inventory available for France, and the ANIS-E database (version v.1.0.1, Violet et al., 2026), which is compiling all NIS reported in European seas from 1492 to 2023, based on available NIS curated databases, such as the Joint Research Centre Data Catalogue for NIS (https://data.jrc.ec.europa.eu/collection/id-00358) used in reports for the European Marine Strategy Directive (NIS descriptor D2). Within the NIS dataset, each species was assigned a biogeographic attribute: i) “NIS-MED”, for species already reported as NIS in the Mediterranean Sea; ii) “NIS-EU”, for species reported as NIS in European seas but not native to or reported as NIS in the Mediterranean Sea; and iii) “Cryptogenic”, *sensu* Carlton (1996), for species whose native or introduced status cannot be confidently determined. These attributes were used to characterize individual species, and all three were considered collectively as NIS in subsequent analyses.

Species not found in the NIS inventories and database were further considered as native species to the Mediterranean Sea (NATS) only if, in the literature, their native range included this region or if they were described as cosmopolitan.

All remaining species were referred to as “uncertain”. This group includes species for which information on their native range is unclear or unavailable, as well as species that are not native to the Mediterranean Sea and have not been reported as NIS in Europe. We did not classify the latter as potential novel NIS (i.e., not yet reported) due to the risk of false-positive identification when using metabarcoding alone (Couton et al., 2022). The complete list of references used to generate these species categories is reported in the “Species occurrence table” (available at the Github repository reported in the Data availability statement).

### Functional traits classification

Functional traits can help elucidate the ecological processes that determine the presence of certain species in specific environments (Mayer-Pinto et al., 2018) or those associated with biological invasions (Quell et al. 2021; Leclerc et al., 2023). Hence, we characterized species using two functional traits, “motility” and “ecology”, selected because they respectively provide information about the dispersal potential of NIS (mobile NIS have greater dispersal potential than sessile ones) and the filtering effect of ports (ports can act as positive filters on species colonizing surfaces and inhabiting the benthos). Motility of species was coded as “sessile” and “mobile”. Ecology was coded for fish using Fishbase (Froese & Pauly, 2000), following Rey et al. (2023) with slight modifications to get a more even distribution, as follows: “demersal” and “bathydemersal” species were grouped into “demersal”, and “pelagic-neritic”, “pelagic-oceanic” and “bathypelagic” species into “pelagic”; “benthopelagic” and “reef-associated” classes were left unmodified. For other species not classified as fish, the modalities for the “ecology” trait were retrieved from the Functional group section available in WoRMS (WoRMS Editorial Board, 2025): species classified as “benthos” and “benthos> macrobenthos” at adult stage were reported as “benthos”; species classified as “plankton”, “plankton>zooplankton” and “nekton” (only *Tursiops truncatus*) at adult stage were classified as “pelagic-non-fish”. An additional trait was considered for fishes, namely “Fishing vulnerability”, which could be related to the greater occurrence of fishes in ports, because of fishing restriction in ports and presence of fishing discards (from recreational fishing boats moored in the port). Vulnerability scores were retrieved from FishBase, where this trait is expressed as a numerical value from 0 (low vulnerability) to 100 (high vulnerability).

### Diversity analyses

Diversity analyses were implemented in R studio using the R package stats (version 4.6.1; R Core Team, 2026) unless otherwise stated in the text. With presence-absence data, alpha-diversity was estimated by species richness. This metric was computed for the whole community (COMM), and separately for the native species (NATS), Non-Indigenous species (NIS, including NIS-MED, NIS-EU and cryptogenic species) and for those species not included in either of the former categories (Uncertain).

The hypotheses of higher NIS numbers (H3) but lower species richness (H4) in ports were tested by generalized linear mixed model (GLMM) performed by using R package glmmTMB (version 1.1.14, Brooks et al., 2017). Habitat type was treated as explanatory variable of species richness (separately for COMM, NATS, NIS and Uncertain dataset) and NIS proportion in the community. Negative binomial and binomial distributions were used to model species richness and NIS proportion in the community, respectively. Model residuals were inspected for overdispersion using the R package DHARMa (Harting, 2026). To take into account our paired sampling (marina and nearby natural habitat) per locality, the locality where samples were collected was included in the model as random factor. To further explore the influence of habitat on species richness and NIS proportion in the community within localities, H3 and H4 were also tested separately in each locality by performing Student’s t-tests or Wilcoxon rank-sum tests when data did not comply the normality or the variance homogeneity assumptions. Finally, differences in species richness and NIS proportion in the community between localities were assessed using one-way ANOVA or Kruskal–Wallis tests when data did not meet the assumptions of parametric tests.

The Jaccard dissimilarity was calculated for COMM, NATS, NIS and Uncertain datasets by the *distance*() function of the phyloseq R package (McMurdy & Holmes, 2013), and used to perform beta-diversity analyses (testing H1 and H2). Principal Coordinates Analyses (PCoA) were used to visualize community’s dissimilarities in an unconstrained ordination space (*ordinate*() and *plot.ordination*() functions of the phyloseq R package). The possibility that communities from ports and natural habitats displayed distinct species composition (H1) was tested performing Partial Distance based Redundancy Analysis (dbRDA) on Jaccard dissimilarities with habitat type as explanatory variable (*capscale*() function of the vegan R package; Oksanen et al., 2013); to control for the potential effect of locality, this factor was included in the partial dbRDA as covariate. The statistical significance of this constrained ordination was assessed by an ANOVA-like test (*anova*.*cca*() function of the vegan R package). Additionally, we estimated the turnover and nestedness components of beta diversity using the betapart R package (version 1.6; Baselga et al., 2023) to determine whether differences in species composition between habitats were primarily driven by species replacement (i.e., turnover) or species loss (i.e., nestedness). Community compositional dispersion was assessed using a permutational analysis of multivariate dispersions (PERMDISP), which was implemented through the *betadisper*() function of the vegan package in R (Oksanen et al., 2013) and was based on Jaccard dissimilarities. This analysis quantifies the dispersion of samples around their habitat-specific centroid in multivariate space. Differences in multivariate dispersion between marinas and natural habitats were tested using the permutation-based test of homogeneity of multivariate dispersions (*permutest*() function in the vegan R package) applied to the *betadisper*() model. The hypothesis of greater biotic homogenization in ports (H2) was considered supported if communities in marinas exhibited significantly lower multivariate dispersion than natural communities, indicating greater similarity in species composition among ports. To investigate whether species detected in different habitats were characterized by different taxonomical and functional traits (H5), species significantly associated with either type of habitat were identified by Indicator Species Analysis using the R package indicspecies (De Cáceres & Legendre, 2009). In addition, species scores on the first constrained axis of the partial dbRDA were used as a measure of species association with the habitat. Differences in the species scores among taxonomic traits (e.g. phylum and class), functional traits (e.g. motility and ecology), and categories (e.g. NATS, NIS, and uncertain) were assessed using a Wilcoxon rank-sum test for two-level traits and a Kruskal–Wallis test otherwise - as the data did not meet the assumptions required for parametric testsu. In case of significant Kruskal-Wallis results, Dunn post-hoc tests with Benjamini-Hochberg “*bh*” correction to account for multiple testing were implemented, for pairwise comparisons between trait levels. For fish species, Spearman rank correlation was performed to assess the relationship between species scores along the first partial dbRDA axis and species-specific fishing vulnerability values.

## Results

### Combining markers with eDNA metabarcoding is an effective way to identify animal taxa across diverse phyla

The outcome of the bioinformatics pipeline analysis is provided in the Appendix S1.2. Each of the three markers revealed a unique set of species (Appendix S1.3-Figure S1, panel A). This was expected with 12S, which has been specifically designed to target teleost species, and is also true for COI and 18S that are known to have complementary affinities. Using three markers also allowed to include species from taxonomical groups which would have been underrepresented by using only one marker (Appendix S1.3-Figure S1, panel B): 12S identified 52.6% of all Vertebrata species, 18S was useful to recover most of Tunicata species (80%) and it was the only marker capable of identifying species from the phyla Xenacoelomorpha, Entoprocta and Placozoa. Similarly, 64.1% of all Arthropoda reported in this study were identified only by COI, which was also the only marker able to detect Chaetognatha. Only one species (i.e., *Dicentrarchus labrax*, belonging to Vertebrata) was detected by all three markers. No other species were common to 12S and 18S. However, 53 species were common to 12S and COI, and 43 species were common to 18S and COI. Combining the taxonomic assignments made with the three markers independently yielded a total of 536 species (see “Species occurrence table” available at the Github repository reported in the Data availability statement), belonging to 18 phyla (Appendix S1.3-Figure S1, panel B). Most of them were among Vertebrata (n=152), followed by Mollusca (n= 89), Annelida (n=81) and Arthropoda (n=54). On average, 125 ± 22 (mean ± SE) species were recorded in each locality.

Of the 536 species (i.e., the entire community, COMM), 354 were native species to the Mediterranean Sea (NATS; Figure 1B), while 132 species were categorized as “Uncertain”, as their native range was either unknown or outside the Mediterranean Sea, and not reported as NIS in Europe (Appendix S.1.4-Figure S2). The remaining 50 species were considered as NIS because they had previously been previously reported as non-indigenous or cryptogenic in Europe (Figure 2). Among these, 14 were classified as “cryptogenic” and seven as “NIS-EU” (i.e., reported as NIS in European seas but not yet in the Mediterranean Sea). The average percentage of NIS per locality (regardless of habitat type) was 6.85 ± 0.6% (mean± SE), reaching up to 13.9 ± 1.71% and 10.3 ± 2.47 % in Sete_1 and Sete_2, respectively (Appendix S1.5: Figure S3). This contribution was not significantly different across the 12 localities (Kruskal-Wallis test, P value = 0.07).

**Figure 2:**
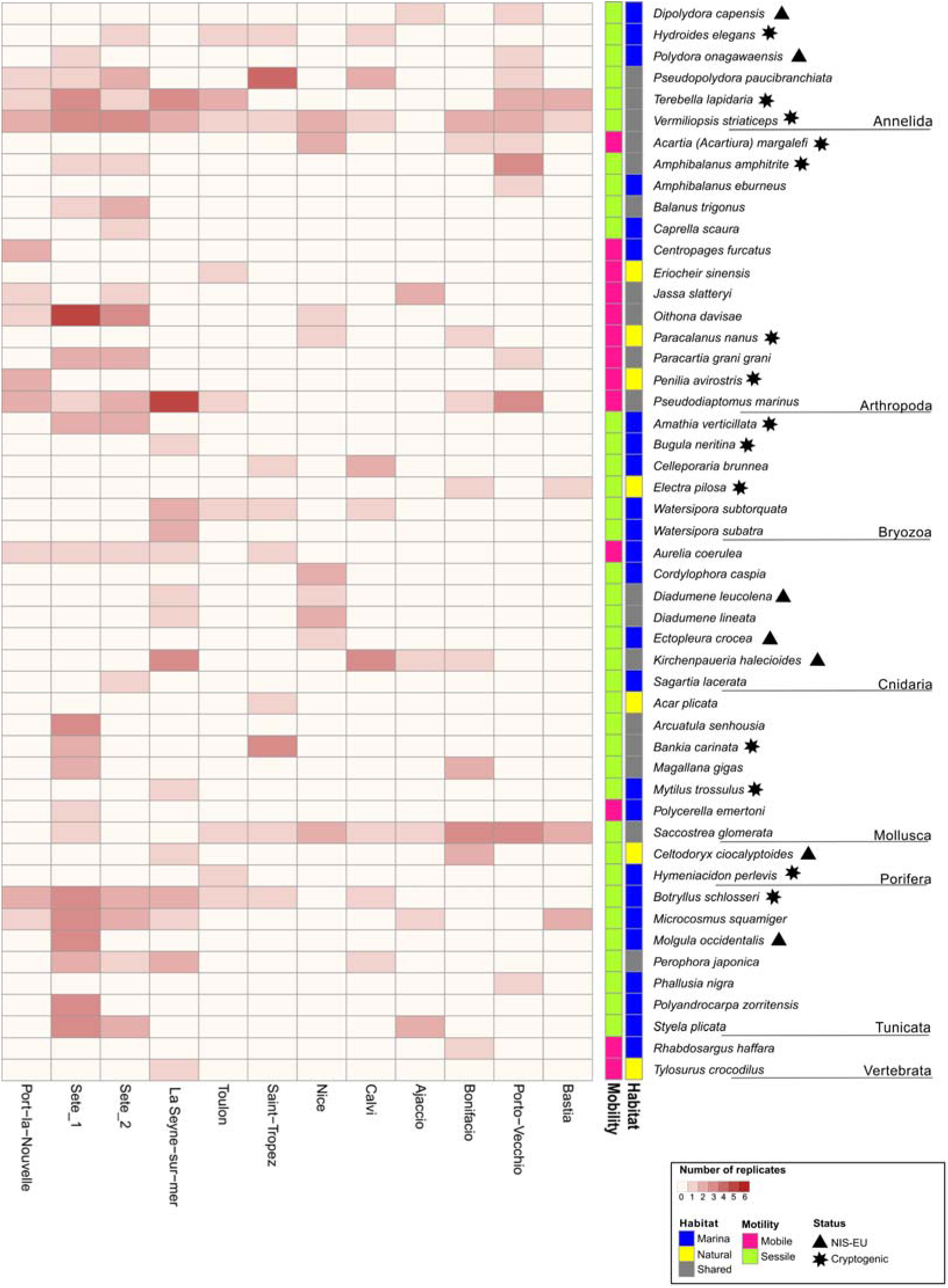
Presence and attributes of NIS (n=50) across localities. The heatmap displays the number of replicates (out of six, with three inside marinas and three in natural habitats) in which each NIS was recorded. Coloured bars on the right display the habitat and motility traits for each species. Symbols highlight species classification as cryptogenic (star) and/or as NIS-EU (triangle), the latter corresponding to species previously reported as NIS in other European seas but neither as native or as NIS in the Mediterranean Sea.

#### Marinas display distinctive and homogeneous animal communities at a regional scale

Across the study region, the animal communities in marinas differed from those in natural habitats, as shown by both unconstrained (Appendix S1.6-Figure S4) and constrained ordinations (Figure 3A). This pattern was statistically supported by a permutation ANOVA (ANOVA.cca) on partial dbRDA, which revealed a significant effect of habitat type on community composition (F-statistic=4.26, p-value = 0.001; R²-adjusted=4.66%), thus supporting H1. Partitioning of Jaccard dissimilarity indicated that the differences between the two habitats were mainly due to species replacement (turnover) rather than to richness-related differences in species composition (nestedness) (Figure 3B). Furthermore, marinas showed significantly lower multivariate dispersion than natural habitats (*PERMDISP*, p-value = 0.006; Figure 3C), indicating greater similarity in species community composition among marinas. This finding supports the hypothesis of increased biotic homogenization in ports compared to in natural habitats (H2). Testing H1 and H2 separately for NATS and NIS revealed similar responses of the two categories of species assemblages to habitat type. Species assemblages differed significantly between marinas and natural habitats for both NATS and NIS (Appendix S1.6-Figure S4). This pattern was confirmed statistically by the ANOVA.cca on partial dbRDA (NATS: F-statistic=4.39, p-value = 0.001, R²-adjusted=4.85%; NIS: F-statistic=3.39, p-value 0.001; R²-adjusted=5.14%; Appendix S1.6-Figure.S5A). This supports H1 for both categories. Again, dissimilarity in both NATS and NIS was primarily driven by turnover, although nestedness contributed more strongly to NIS than to NATS, accounting for 15.4 ± 4.7% and 7.7 ± 2.6%, respectively, of the total Jaccard dissimilarity (Appendix S1.6, Figure S5B). Likewise, both NATS and NIS displayed biotic homogenization in marinas (H2) (*PERMDISP*; NATS: F-statistics=7.39, p-value = 0.004; NIS: F-statistic=10.7, p-value =0.002; Appendix S1.6-Figure S5C).

**Figure 3:**
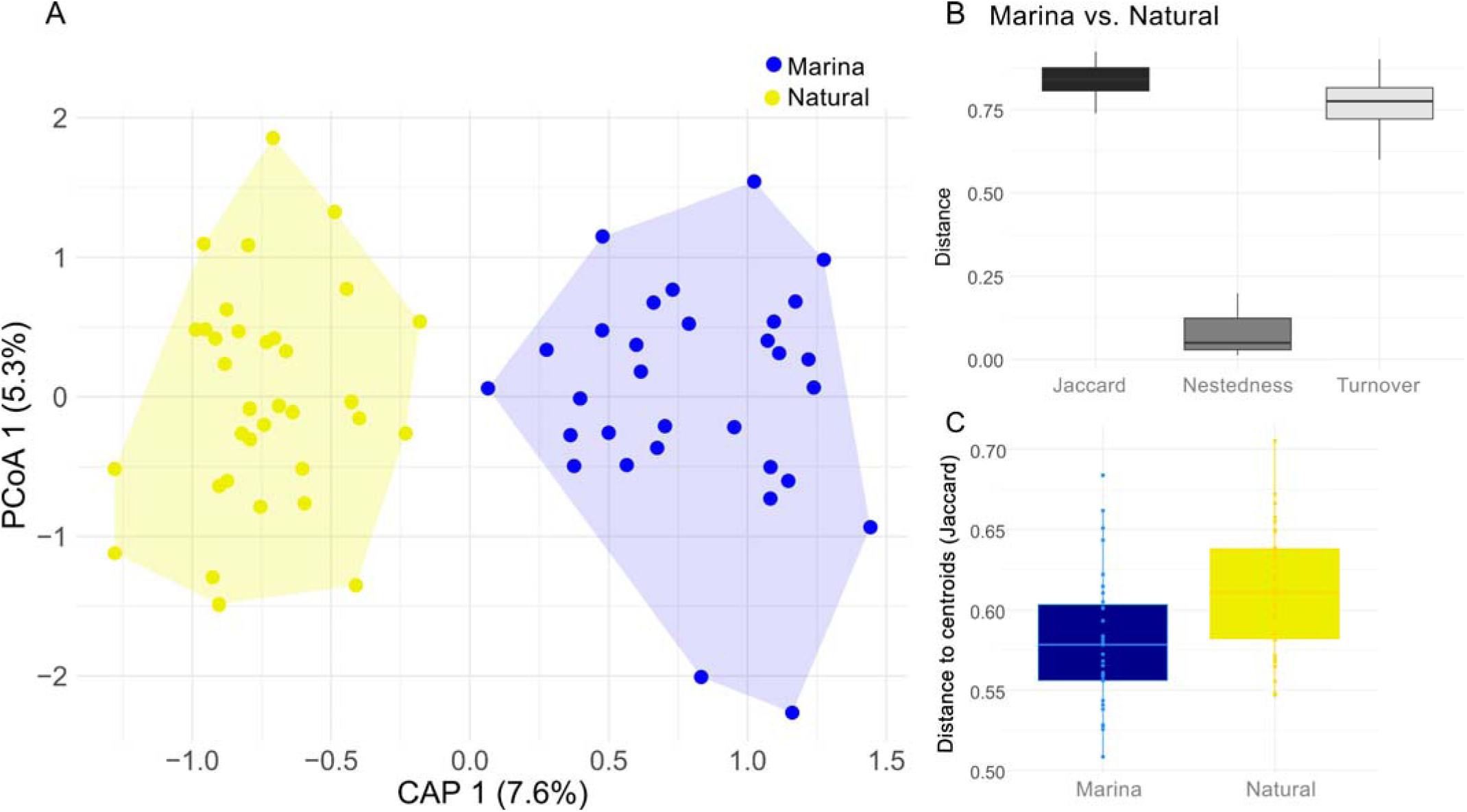
Jaccard dissimilarities based on species composition of communities (COMM) in marinas andin natural habitats. A) Partial dbRDA showing the effect of habitat on community while accounting for the locality. Because only one explanatory variable was included, the analysis yielded a single constrained axis (CAP 1) and one unconstrained axis (PCoA 1); B) boxplots displaying partitioning of Jaccard dissimilarities into nestedness and turnover components, when comparing samples from inside marinas and natural sites; C) boxplot showing dispersion of Jaccard dissimilarities among samples collected inside marinas (blue) and natural habitats (yellow).

Finally, H1 was also statistically supported for the Uncertain group (ANOVA.cca: F-statistic=2.63, p-value 0.001; R²-adjusted=2.36%; Appendix S1.6-Figure S5A). As observed for the other categories, species assemblages’ dissimilarities between the two habitats were mainly driven by turnover (Appendix S1.6-Figure S5B). Conversely, this category did not show evidence of biotic homogenization in marinas (*PERMDISP*, F-statistics= 0.99, p-value = 0.33) (Appendix S1.6-Figure S5C).

#### NIS are overrepresented in marinas compared to natural habitats

In line with our working hypothesis (H3), the proportion of NIS in marinas (10.4 ± 0.8%, mean ± SE) was almost three times that in natural habitats (3.5 ± 0.6%). This difference was statistically supported by the results of the GLMM (natural habitat relative to port: β= -1.044, 95% CI [-1.38; -0.70], SE=0.17, z value = -5.95, p-value < 0.0001). With the exception of Bonifacio (Figure 4), this trend was observed in every locality and significant in seven of them, namely Bastia, Calvi, Nice, Port-la-Nouvelle, Porto-Vecchio, Sete_1 and La Seyne-sur-Mer (Student’s t-test and Wilcoxon’s test results available in Appendix S2-Table S3).

**Figure 4:**
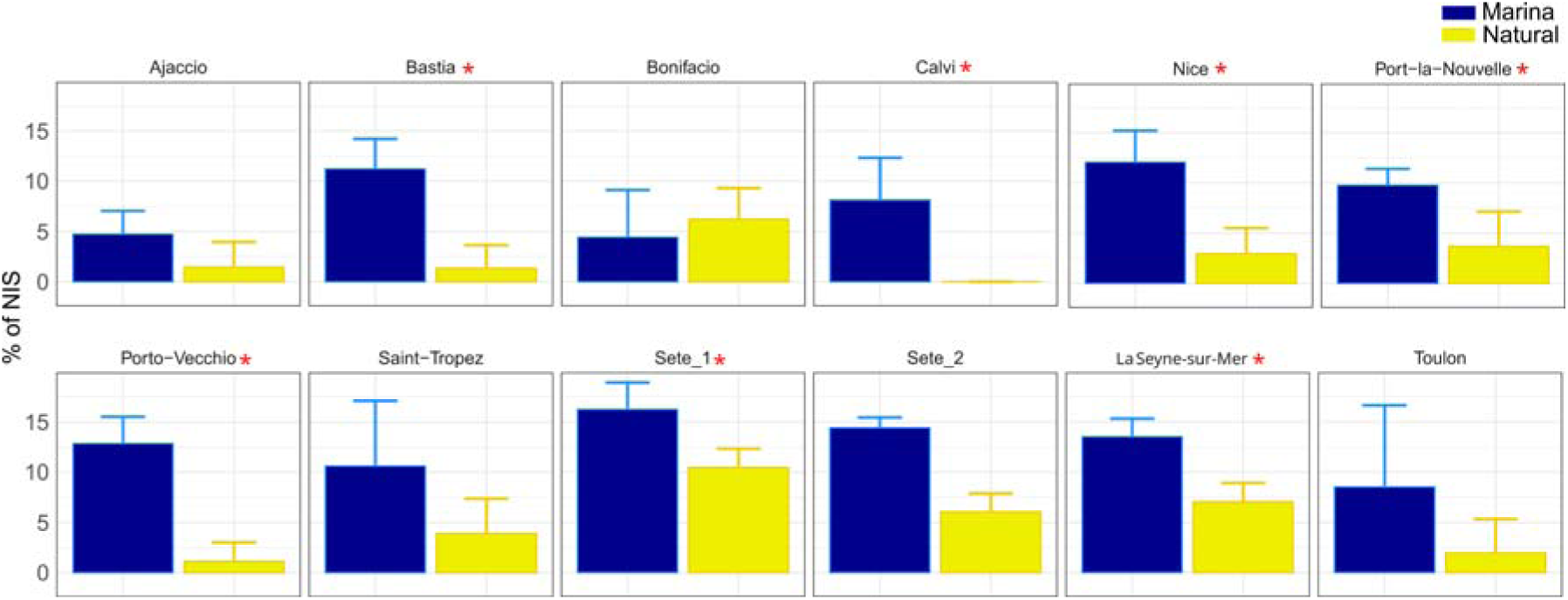
Percentage of NIS. Barplot showing mean and standard error of the percentage of NIS in communities sampled in marinas and in natural habitats in each locality. Localities that showed a significant difference in the proportion of NIS inside and outside marinas are labelled with * (Student’s t-test or Wilcoxon test, one-tailed, p-value < 0.05).

The number of NIS was also significantly higher in marinas across all localities (GLMM, natural habitat relative to port: β= -1.41, 95% CI [-1.74; -1.09], SE=0.16, p-value < 0.0001) (Figure 5 and Appendix S1.8-Table S1) and per locality (Appendix S1.7-Figure S6). In addition, nearly five times more NIS were observed exclusively in marinas (25 NIS) than in natural habitats (7 NIS) (Figure 5). It is however worth noting that ca. 43% of NIS detected in marinas were also present in natural habitats (18 NIS; Figure 5).

**Figure 5:**
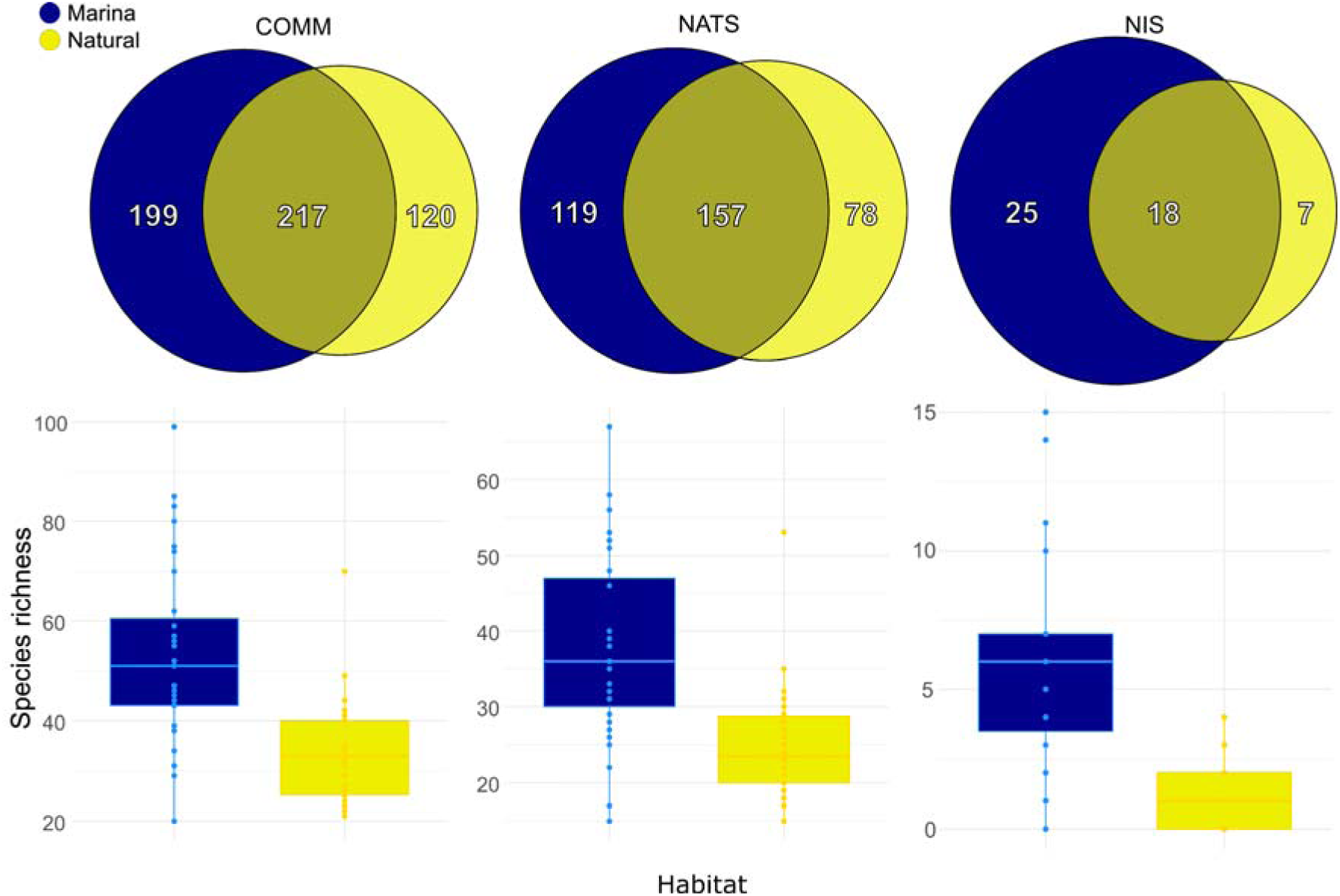
Number of species private and shared between habitats. Venn diagrams showing the total number of species exclusive or shared in marinas and natural habitats, considering the whole community (CO M), non-indigenous species (NIS) only and native species (NATS) only. Boxplots of species richness per sample calculated for each habitat type are also provided.

The greatest NIS richness was found in the two study localities of Sete, both inside and outside marinas (Sete_1: average 13.3± 2.1 NIS inside and 3.5 ± 0.7 NIS outside; Sete_2: average 12.5 ± 3.5 NIS inside and 2.5 ± 0.7 NIS outside).

#### Marinas host richer communities regardless of high NIS occurrence

Contrary to our working hypothesis (H4), the studied marinas did not exhibit lower species richness than nearby natural habitats (COMM dataset, Figure 5). On the contrary, species richness was significantly higher in marinas overall (GLMM, natural habitat relative to marina: β= -0.45, 95% CI [-0.58; -0.32], SE=0.06, z value= -6.88, p-value < 0.0001; Appendix S1.8-Table S1). This pattern was consistently observed across localities, with the exception of Bonifacio (Appendix S1.8-Figure S7). Significant differences were detected in Calvi, La Seyne-sur-Mer and Porto-Vecchio (Appendix S1.8-Table S2). Considering phyla separately, H4 was supported only for Arthropoda (GLMM, natural habitat relative to port: β= 0.35, 95% CI [0.11; 0.61], SE=0.12, z value= 2.85, p-value = 0.004; Appendix S1.8-Table S3). In contrast, nine phyla displayed greater richness in marinas than in natural habitats: Mollusca, Cnidaria, Vertebrata, Bryozoa, Annelida, Tunicata, Porifera, Nemertea and Rotifera (Appendix S1.8-Table S3 and Figure S7), while the eight remaining phyla did not show significant differences.

The higher richness observed in marinas was not driven only by the higher NIS occurrence within them: testing H4 on NATS and Uncertain dataset alone produced similar results of higher richness in marinas than nearby natural habitats (Figure 5; GLMM for NATS: natural habitat relative to marina: β= -0.40, 95% CI [-0.54; -0.26], SE=0.07, z value = -5.68, p-value < 0.0001; GLMM for Uncertain; natural habitat relative to marina: β= -0.34, 95% CI [-0.51; -0.61], SE=0.09, z value = -3.75, p-value = 0.0001). When examining each locality separately, significant differences between habitats were observed only for NATS in Calvi and for Uncertain in Sete_1, with higher richness in marinas than in nearby natural habitats (Student’s t-test, p-value = 0.01, results of tests on all localities reported in Appendix S2-Table S4).

#### Taxonomical and functional traits drive differential species occurrence in marinas and natural habitats

Community differences between marinas and natural habitats were associated with the differential occurrence of 60 species (including species from the NATS, NIS and Uncertain categories) identified by Indicator Species Analysis (IndVal.g statistic, p-value < 0.05, Appendix S1.9-Table S4; Figure 6A). A substantially larger number of species were identified as indicators of marinas (53 species) than of natural habitats (7 species), suggesting again a distinct community in the former. Most indicator species were Teleostei in both habitats (33.9% of indicator species of marinas and 42.8% of natural habitats), followed by Polychaeta in marinas (18.8%) and Copepoda (42.8%) in natural habitats. NIS accounted for 16% of marinas indicator species while none was significantly associated with natural habitats (Figure 6A).

**Figure 6:**
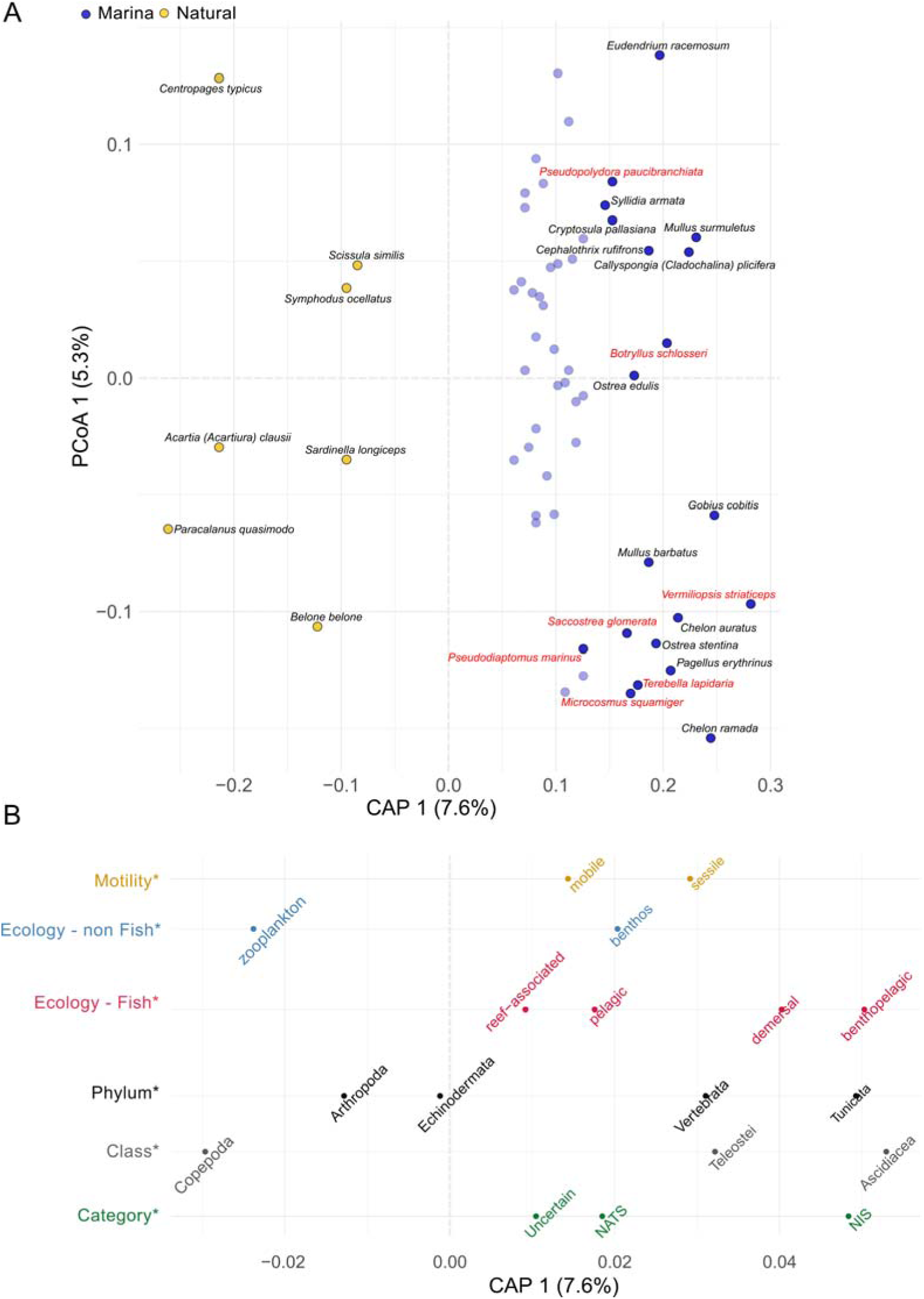
Species scores of the partial dbRDA performed to test the effect of habitat on COMM species composition accounting for locality; A) species scores of the 60 indicator species identified for each habitat by Indicator Species Analysis: species labels are displayed for all 7 indicator species of natural habitats (yellow) and only for the 20 indicator species of marinas (blue) displaying the highest statistics in the permutation test (Indval.g); NIS are reported in red; B) average values of species scores of partial dbRDA first axis obtained for each functional and taxonomic group and for the origin category in which they were included (i.e. NIS, NATS or Uncertain): traits that significantly influenced the distribution of species scores in the first dbRDA axis are flagged with an asterisk (for tests performed, statistics and p-values see Appendix S1.9-Table S5). Values located towards the right side of the first dbRDA axis are more strongly associated with marinas whereas those towards the left side were associated with natural habitats.

The distribution of species scores of the COMM dataset along the first axis of the partial dbRDA was analysed to determine if differences in species occurrence between habitats were related to taxonomic or functional traits (Figure 6B). Species scores differed significantly among Phyla (Kruskal-Wallis test, p-value < 0.001), Classes (Kruskal-Wallis, p-value = 0.01), motility trait (Wilcoxon signed-rank test, P value = 0.002), ecology trait (for non-fish species, Wilcoxon rank-sum test, p-value < 0.001; for fish species, Kruskal-Wallis test, P value = 0.04) and category (NATS, NIS, Uncertain) (Kruskal-Wallis test, P value = 0.002) (Appendix S1.9-Table S5). In contrast, the distribution of fish species scores in the two habitats did not appear to be correlated with their level of vulnerability to fishing (Spearman correlation between fish species scores on the dbRDA first axis and fishing vulnerability values, p–value = 0.07).

As reported in Figure 6B and Appendix S1.9-Table S5, Vertebrata and Tunicata tended to be associated with marinas (i.e., species scores displaying positive values of dbRDA axis), and Arthropoda and Echinodermata with natural habitats (i.e., negative values of dbRDA axis) (Post-hoc Dunn tests, p-value < 0.05). At Class level, species in marinas were mostly assigned to Teleostei and Ascidiacea while those found outside were mainly assigned to Copepoda. Marinas also had a higher occurrence of sessile and benthic non-fish species, as well as demersal and benthopelagic fishes. In contrast, mobile species, pelagic-non-fish and fishes classified as “reef-associated” were more frequently recorded in natural habitats than in marinas. Finally, species scores were significantly greater for NIS than for NATS and Uncertain categories (post hoc Dunn tests, p < 0.01), indicating a stronger association of NIS with marinas.

Testing the influence of traits on NATS dissimilarities yielded results broadly similar to those obtained for COMM, but the distribution of species from this assemblage in the two habitats did not appear to be associated with the motility trait or with the ecology trait for fishes (Appendix S1.9-Table S5). Lastly, except for a marginally significant influence of “phylum” on NIS (Kruskal-Wallis, p-value=0.04), no significant effect of taxonomic and functional traits was observed either for NIS or for the Uncertain species datasets.

## Discussion

### What explains higher species richness in marinas, across phyla?

By combining the taxonomic assignment made with three markers, our eDNA metabarcoding approach allowed the identification of 536 animal species across 18 phyla, in 65 seawater samples collected in 12 pairs of recreational marinas and adjacent natural habitats along the French Mediterranean coast. These included 50 species classified as non-indigenous or cryptogenic based on previous records. Our study revealed marked differences between marinas and natural habitats in community richness, structure and functional traits. Beyond the expected higher prevalence of NIS in ports, a less anticipated result (contrary to our hypothesis H4) was the overall greater taxonomic richness in the study marinas than in the adjacent natural habitats, including when considering native species alone. This pattern was observed across multiple phyla, which encompass a diversity of life-history traits, with significantly higher richness in nine, no significant difference in eight and significant lower richness in only one, namely Arthropoda. On the one hand, our finding is consistent with previous studies reporting higher taxonomic richness in artificial habitats, including marinas and ports, for particular taxonomic groups, including Polychaeta (Dafforn et al., 2013), Foraminifera and benthic macrofauna (Pavard et al., 2026), and fishes (Macé et al. 2026; Ntouni et al., 2023). On the other hand, several studies have supported the view that species diversity is generally lower in artificial coastal habitats than in adjacent natural habitats (e.g. Wakefield et al., 2013; Bulleri & Chapman, 2010; Bishop et al., 2022). The higher richness observed across multiple phyla in our study therefore calls for a closer examination of potential methodological and ecological explanations.

First, methodological aspects deserve attention, as species presence was inferred from eDNA signals rather than direct observations of organisms. Although the study ports were all marinas – hosting exclusively recreational boats and thus not subject to ballast water discharges – they are continuously exposed to the introduction of biological material transported as biofouling on the hulls of resident or visiting recreational boats (Ulman et al., 2019; Ashton et al. 2022). Thus, the greater community richness observed in this study may reflect the detection of DNA from resident species alongside DNA transiently introduced by boats from outside the marina. Although this scenario is plausible, the eDNA signal from transient species is expected to be weaker and more short-lived than that of resident species, resulting in fewer sequencing reads. Furthermore, we performed rarefaction of the raw sequencing data. This subsampling of the original reads to a fixed sequencing depth further reduced the probability of detecting low abundance taxa, including rare and transient species (Shirazi et al. 2021). Another methodological aspect to consider is the distinct hydrodynamic conditions of ports relative to adjacent natural habitats. The higher richness observed in marinas could potentially result from longer water residence times within ports compared to natural habitats. This mechanism may be particularly relevant in enclosed harbour systems, such as many large commercial ports. However, it is less likely to explain the patterns observed here, as all studied marinas were directly connected to the sea and lacked gates, locks, or other barriers that restrict water exchange. Finally, similar patterns of higher taxonomic richness in artificial than natural habitats have also been reported in studies based on direct observation (i.e. sampling or visual survey of specimens) rather than eDNA metabarcoding, including studies on Polychaeta (Dafforn et al., 2013), Foraminifera and benthic macrofauna (Pavard et al., (2026), and fishes (Ntouni et al., 2023). Taken together, these observations suggest that methodological biases associated with the transient introduction and transport of eDNA are unlikely to fully explain the greater richness detected in marinas.

Several ecological and environmental factors may also contribute to the higher species richness observed in marinas. Among these, the heterogeneous mosaic of substrates found in ports deserves particular attention. Ports, from commercial harbours to marinas, are infrastructures often replacing or modifying pre-existing soft-sediment habitats, and introducing a diversity of artificial materials and structural complexity that creates a wide range of microhabitats, suitable to support a diversity of species assemblages (Connell & Glasby, 1999; Todd et al., 2019; Romoth et al., 2023). In addition, nutrient enrichment in ports (Sebastiá & Rodilla, 2013; Guerra-García et al., 2021) may enhance primary production (Scott, 1978), and, under certain conditions, secondary production (Leslie et al., 2005), favouring taxonomic diversity (Whittaker et al., 2003). Port regulations may also contribute to higher species richness, although their direct effects are likely to be restricted to particular taxonomic groups, notably fishes and other harvested species. Fishing restrictions in marinas may provide favourable conditions for fish populations, notably by enhancing their use as foraging and nursery grounds. Fish communities have indeed repeatedly been shown to be as diverse –or even richer-than those in natural habitats (e.g., Bouchoucha et al., 2016; Manel et al., 2024; Macé et al., 2024, 2026). In addition, greater fish presence may also exert top-down control on space-dominant competitors among epibiota, enhancing community diversity (Rivero et al., 2013). Finally, intermediate levels of disturbance, common in small ports such as the marinas considered in this study, may further promote diversity by enabling the coexistence of both subordinate and dominant species (Dial & Roughgarden, 1998; Santos et al., 2021).

Overall, the greater community richness observed in marinas may reflect the combined influence of methodological (transient eDNA), structural (habitat heterogeneity), regulatory (fishing restrictions) and ecological (productivity, species interaction etc.) factors. Further studies based on in-situ observation or targeting substrate-specific assemblages across different habitats, are needed to disentangle the relative contribution of these factors.

### Similarities and distinctiveness of marinas: the ecological dimension of the “portuarization syndrome”

At the regional scale, we found greater similarity among marina communities than among natural habitats, supporting the hypothesis of biotic homogenization. This finding also suggests that the concept of “biological portuarization syndrome”, which was proposed by Touchard et al. (2023) to describe repeated port-induced evolution across broad spatial scale, could be extended to include an ecological dimension. This homogenization may be the result of two complementary processes: the environmental filtering of autochthonous species (Todd et al., 2019) and the high connectivity of ports through boating, which uniformizes species assemblages across scales (Touchard et al., 2023). This finding has recently been reported for fishes’ communities (Macé et al. 2026) but, to our knowledge, out study is the first evidence for biotic homogenization in ports across multiple phyla at a regional scale.

The portuarization syndrome is also evident through species assemblages in ports that differ significantly from those in nearby natural areas. As we used the same standardized sampling and analytical protocol for every sampling locality, this observation remains valid regardless of the species name obtained through the taxonomic assignment procedure (i.e., molecular operational taxonomic units can be compared between marinas and natural habitats, regardless of their assigned taxonomic names). Interestingly, this dissimilarity between habitats was not primarily due simply to species losses (nestedness) between natural sites and marinas, but rather to extensive species turnover. This pattern scales up to the Class level, with Teleostei and Copepoda exhibiting contrasting associations with the two habitats, as revealed by the species scores of partial dbRDA. One possible explanation for this observation is that zooplankton populations are strongly regulated by fish larvae and adults (Gliwicz & Pijanowska, 1989). Higher fish densities in marinas could reduce copepod species richness, through species-specific and size-dependent selective predation (Fiksen et al., 2005). Moreover, zooplankton are highly sensitive to water temperature, transparency and oxygen concentration (Stoch et al., 2023), which most often differ between ports and adjacent natural habitats. This may further influence the composition and occurrence of zooplankton populations.

Another important factor distinguishing communities in marinas from those in natural sites was linked to the studied functional traits, namely species motility and ecology. Functional trait analyses confirmed the expectation of a rich and abundant fouling community in ports (Tempesti et al., 2022), characterized by sessile and benthic species. The higher association of demersal and benthopelagic fishes than pelagic ones with marinas was also expected given the reliance of the former species on benthic preys (Bergstad, 2019). By contrast, reef-associated fish were less common in marinas. This may be because, despite the variety of artificial substrates, these habitats do not necessarily provide the same reef-like structural features as natural rocky habitats, such as crevices.

Lastly, the assemblages in marinas were characterized by a greater occurrence of NIS compared with the other two species categories (NATS and Uncertain). Thirty-six percent of NIS were shared between marinas and natural habitats (18 shared detections across 50 NIS species), which suggests either restricted spreading from one habitat to the other, or the parallel introduction of these NIS in the two habitats. A further 14% of NIS were only detected outside marinas, likely reflecting the existence of alternative introduction vectors besides shipping and boating – such as aquaculture, a particularly relevant vector in France (Massé et al., 2023). Beyond a weak effect of phylum, none of the traits studied provided clear insights into the habitats in which NIS were more likely to occur. Although trait-based analyses remain challenging because many NIS are still poorly studied (but see e.g., Quell et al., 2021 and Leclerc et al., 2023), future research should expand beyond mobility and broad ecological traits to consider characteristics directly linked to invasion success, including tolerance to environmental stressors and pollutants, feeding strategies, dispersal traits and growth rates. Such information may improve our ability to predict the likelihood of NIS establishment and spread. It is important to note that the functional trait patterns identified in this study should be regarded as exploratory, as they were inferred from the distribution of species scores on the first dbRDA axis. Nevertheless, our observations provide valuable insights into the potential filtering effect of port habitats on species based on functional traits, and may inform further experimental work to test this hypothesis.

### Are NIS overlooked in the wild?

A total of 50 non-indigenous species (NIS) were identified across all localities, including seven that have not yet been recorded in the Mediterranean Sea, but are known as NIS in other European seas. We acknowledge that it is not possible to confirm their presence using eDNA alone (i.e., false positives may occur, see Couton et al., 2022). However, we tried to minimize false assignments as much as possible, for instance during the bioinformatic processing by excluding rare taxa with using a rarefaction procedure, and during the taxonomic assignment step by using thresholds previously tested for their reliability with the markers used (Couton et al. 2019). Confidence in NIS detections made through eDNA metabarcoding also increases when combining the detections from multiple markers (Pappalardo er al., 2026). In our study, nine NIS were detected by both 18S and COI, namely *Amphibalanus amphitrite*, *Kirchenpaueria halecioides*, *Magallana gigas*, *Penilia avirostris*, *Pseudodiaptomus marinus*, *Pseudopolydora paucibranchiata*, *Styela plicata*, *Terebella lapidaria* and *Watersipora subtorquata*. We also carefully checked the assignment and reference data for particularly confusing taxa, such as *Watersipora subtorquata* vs. *W. subatra*, (Gauff et al. 2023a), and *Botrylloides* spp., (Viard et al. 2016). Lastly, another criterion that increases the validity of NIS eDNA metabarcoding detections, is “biological plausibility”: the likelihood that a species will be found in a specific environment or geographic location (Pappalardo et al., 2026). Several non-indigenous species (NIS) that were detected in this study through eDNA metabarcoding were previously reported in Mediterranean ports through large-scale NIS inventories based on in situ collection and morphological identification (Ulman et al., 2017; Tempesti et al., 2020; Galanidi et al., 2023). Among the taxa that we detected, the most frequently reported species in these inventories included representatives of several phyla: Annelida, such as *Hydroides elegans and Pseudopolydora paucibranchiata*; Arthropoda, including *Amphibalanus eburneus*, *Balanus trigonus*, *Caprella scaura*, *Eriocheir sinensis*, *Oithona davisae*, *Paracartia grani grani*, and *Pseudodiaptomus marinus*; Bryozoa, represented by *Amathia verticillata* and *Celleporaria brunnea*; Cnidaria, represented by *Diadumene lineata*; and Tunicata, including *Microcosmus squamiger*, *Phallusia nigra*, *Polyandrocarpa zorritensis*, and *Styela plicata*. Furthermore, some of our detections were confirmed by previous visual observations in the same port locations, which provides independent validation. For instance, the mollusks *Arcuatula senhousia* and *Polycerella emertoni* were previously observed in Sete_1 in 1989 and in 2018, respectively (Massè et al., 2023). Likewise, the Bryozoans *Celleporaria brunnea* and *Watersipora subtorquata* were reported in Saint-Tropez by Ulman et al., (2017) and in Toulon by Gauff et al., (2023b), respectively. Gauff et al. (2023b) also reported the presence of the Annelid *Hydroides elegans* in Toulon. The limited availability of morphology-based inventories conducted specifically in the marinas included in our study limits further comparisons. However, our results help identify localities in which further investigation should be prioritized for targeted molecular investigation and/or surveys based on direct observations. According to our detections, one locality displayed the highest NIS richness and the greatest proportion of NIS in the community: Sete_1. This result was not unexpected, as this site is surrounded by other urban areas with marinas and is next to the Thau lagoon — a well-known introduction hotspot in France. This is due to the large-scale import of Japanese oysters (*Magallana gigas*) in the 1970s (Boudouresque et al., 2010), which frequently carried other species on their shells and on the material used to collect and transport spats. This illustrates the close relationship between the diversity of introduction vectors/pathways and NIS establishment (Boudouresque et al., 2010).

Unsurprisingly, most NIS were detected in marinas, with 25 occurring exclusively in these habitats. By comparison, 18 NIS occurred in natural habitats, of which only seven were exclusive of them. A similar pattern was observed for NIS proportions. The proportion of NIS in marinas, though generally higher, was however not markedly greater than in natural habitats, with only five localities (Bastia, Calvi, Seyne-sur-Mer, Porto-Vecchio, and Nice) showing a significant increase in NIS proportions in marinas. So far, little attention has been given to natural habitats near ports, with NIS being primarily investigated only inside ports (Madon et al., 2023; Zarcero et al., 2024; Zarcero et al., 2026) or in distant and pristine sites such as marine protected areas (Giakoumi & Pey, 2017). Our results suggest that natural habitats adjacent to marinas may play a critical but underappreciated role in the spread of NIS by acting as recipient habitats for spillover from ports. They could then act as stepping-stone locations for the colonization of more distant and pristine habitats. This process may be particularly important if transient establishment in these intermediate habitats allows NIS to acclimate or adapt to local natural conditions. Natural habitats located near marinas are also likely to experience some anthropogenic pressures. These disturbed conditions may lower competitive resistance from native species (Giakoumi & Pey, 2017), making these habitats particularly susceptible to secondary establishment by NIS. Therefore, it is crucial to survey NIS presence in these locations not only because it can inform about NIS capable to spread further but also because these areas represent the majority of coastal zones in the highly urbanized Mediterranean Sea.

Overall, our study revealed that marinas are introduction hotspots harbouring unexpectedly high diversity across animal phyla and homogeneous communities at a regional scale. This provides ecological perspectives to extend the concept of “portuarization syndrome”: a perspective that, in future, should be tested on ports with different dimensions and main use from recreational marinas. The substantial NIS presence in adjacent natural habitats simultaneously underscores the need for expanding monitoring and NIS studies beyond ports to better understand the spatial dynamics of marine bioinvasions and the consequences of the expanding urban sprawl on their natural counterparts in coastal zones.

## Supplementary Information

Appendix_S1.doc – Figures S1-S7; Tables S1-S5

Appendix_S2.xlsx – Tables S1-S4

## Data Availability Statement

The codes and datasets used to perform the analyses, including the OTUs tables generated for each of the three marker genes (12S, 18S and COI), as well as the Species Occurrence Table, are publicly available in the Github repository https://github.com/blind-submissions-2025/Portuarization_MedSea_eDNA.git. The 12S, 18S and COI gene sequence data generated and analysed in this study have been deposited under the BioProject number PRJEB103747. Appendix_S1 and Appendix_S2 cited in the manuscript are available as supporting information.

## Author contributions

**GL**: Methodology (equal); Data curation (lead); Formal analysis (lead); Visualization (lead); Writing – original draft (lead); Writing – review and editing (lead). **ACJ**: Methodology (equal); Investigation (equal); Resources (equal); Formal analysis (supporting); Software (equal); Writing – review and editing (Supporting). **CV**: Data curation (supporting); Resources (supporting); Writing – review and editing (Supporting). **MB**: Methodology (supporting); Resources (equal); Investigation (supporting); Writing – review and editing (Supporting). **XT**: Resources (equal); Formal analysis (supporting); Writing – original draft (supporting); Writing – review and editing (equal). **SAH**: Conceptualization (equal); Methodology (equal); Resources (equal); Data curation (equal); Formal analysis (supporting); Funding acquisition (equal); Writing – original draft (supporting); Writing – review and editing (equal). **FV**: Conceptualization (equal); Methodology (equal); Investigation (supporting); Resources (equal); Formal analysis (supporting); Data curation (supporting); Funding acquisition (equal); Project administration (lead); Writing – original draft (equal); Writing – review and editing (equal).

## Acknowledgements

The authors wish to thank all partners involved in the 2021 SUCHIMED campaign (RV L’Europe, https://doi.org/10.17600/18001619), and especially, Françoise Miralles, Fabienne Chavanon and Christophe Ravel. The authors are grateful to Pierre Taberlet for sharing the unpublished sequences for the primers of the teleo4 marker, to Argyro Zenetos for her thorough review of the NIS list presented in this study and for offering insightful feedback, and to Daniel Martin and Iosune Uriz for providing expert advice on the native range of Annelida and Porifera species. This work benefited from funding through the French National Research Agency (ANR) under the “Investissements d’Avenir” program with the reference ANR-16-IDEX-0006 (i-siteMUSE) supporting the MarEEE project. Data used in this work were partly produced through the GenSeq technical facilities of the « Institut des Sciences de l’Evolution de Montpellier » with the support of LabEx CeMEB, an ANR “Investissements d’avenir” program (ANR-10-LABX-04-01). This research was co-funded by the European Union (GA#101059915) supporting the BIOcean5D project through the funding of GL post-doctoral fellow. Views and opinions expressed are however those of the author(s) only and do not necessarily reflect those of the European Union. Neither the European Union nor the granting authority can be held responsible for them.

