## Appendix_S1 for "Environmental DNA reveals biotic homogenization alongside elevated native and non-native species richness in marinas, supporting an ecological portuarization syndrome"

### Appendix S1.1: Study marinas

Characteristics of the marinas included in this study: number of berths, maximum draft and maximum boat length were retrieved from official port websites. Additionally, information about ports compliance with the ISO 18725 standard about Clean Ports established by UPACA (Union des Ports de Plaisance Provence Alpes Côte d'Azur et Monaco; [*www.ports-propres.org*](http://www.ports-propres.org)) is also reported.

| **Marina** | **Berths** | **Max draft (m)** | **Max boat length (m)** | **Type of lines** | **Certifications "Clean Ports"** |
| --- | --- | --- | --- | --- | --- |
| Port-la-Nouvelle | 334 | 4.5 | 12 | cataway | Clean Port |
| Sete_1 (Quilles) | 377 | 1.5 | 8 | *n.a.* | *n.a.* |
| Sete_2 (Sur-de-France) | 820 | 8 | 50 | Mooring buoys; gang ways | Clean Port |
| La Seyne-sur-Mer | 300 | 3.5 | 14 | Mooring buoys; stern to the dock | Clean Port; Active in Biodiversity |
| Toulon (Vielle Darse) | 560 | 8 | 55 | Mooring buoys; stern to the dock | Clean Port; Active in Biodiversity |
| Saint Tropez | 734 | 3.5 | 18 | Mooring buoys; stern to the dock | Clean Port; Active in Biodiversity |
| Nice (Lympia) | 488 | 4.5 | 18 | Mooring buoys;Catway; Alongside; Stern to the dock | *n.a.* |
| Bonifacio | 350 | 6 | 85 | Mooring buoys; stern to the dock | Clean Port; Active in Biodiversity |
| Bastia (Port Toga) | 357 | 3.2 | 30 | Mooring buoys; stern to the dock | *n.a.* |
| Calvi | 500 | 7 | 65 | Mooring buoys; stern to the dock | *n.a.* |
| Ajaccio (Tino Rossi) | 280 | 12 | 100 | Mooring buoys; stern to the dock; cataway; alongside | Clean Port; Active in Biodiversity |
| Porto Vecchio | 380 | 3.5 | 50 | Mooring buoys; stern to the dock | *n.a.* |

### Appendix S1.2: Outputs of the bioinformatic pipeline

|  | | **COI** | **12S** | **18S** |
| --- | --- | --- | --- | --- |
| **N reads for DADA input** | 37337818 | | 26359265 | 35976645 |
| **N reads before chimera removal** | 32158146 | | 24704831 | 33240665 |
| **N reads after chimera removal** | 26692757 | | 21098423 | 23211105 |
| **N reads post decontamination** | 26692723 | | 19078806 | 23205567 |
| **N reads in final output** | 25993198 | | 18658635 | 22646411 |
| **N ASV in final output** | 55422 | | 5039 | 33530 |
| **N OTU in final output** | 16175 | | 2835 | 22300 |
| **N OTU post rarefaction to 100.000 reads** | 7466 | | 1546 | 10298 |
| **N OTU assigned to Eukaryotes** | 663 | | 682 | 2419 |
| **N OTU assigned to Animalia** | 422 | | 562 | 376 |
| **N OTU assigned to species** | 372 | | 455 | 260 |
| **N species identified** | 290 | | 144 | 200 |

### Appendix S1.3: Complementarity between the three markers used

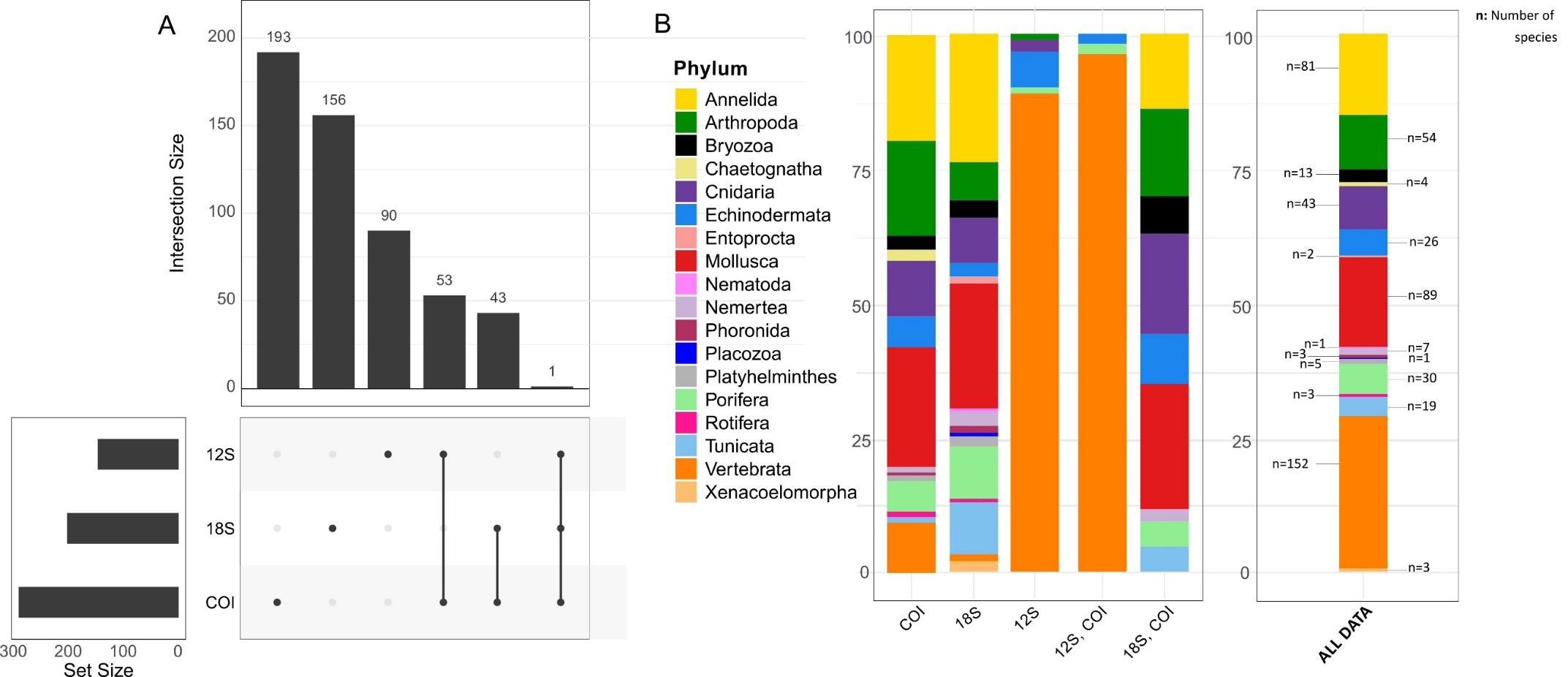

*Figure S1: A) Number of species classified by each marker and by their combination in the Species occurrence table; B) Proportion of species classified by each marker and their combination (i.e. different combinations are reported on the x axis) and assigned to different phyla. The proportion and the number of species assigned to different phyla across the whole dataset (i.e. merged over the three markers and all samples) is reported and labelled “All data”.*

### Appendix S1.4: Composition of the “Uncertain species” category in the different habitats

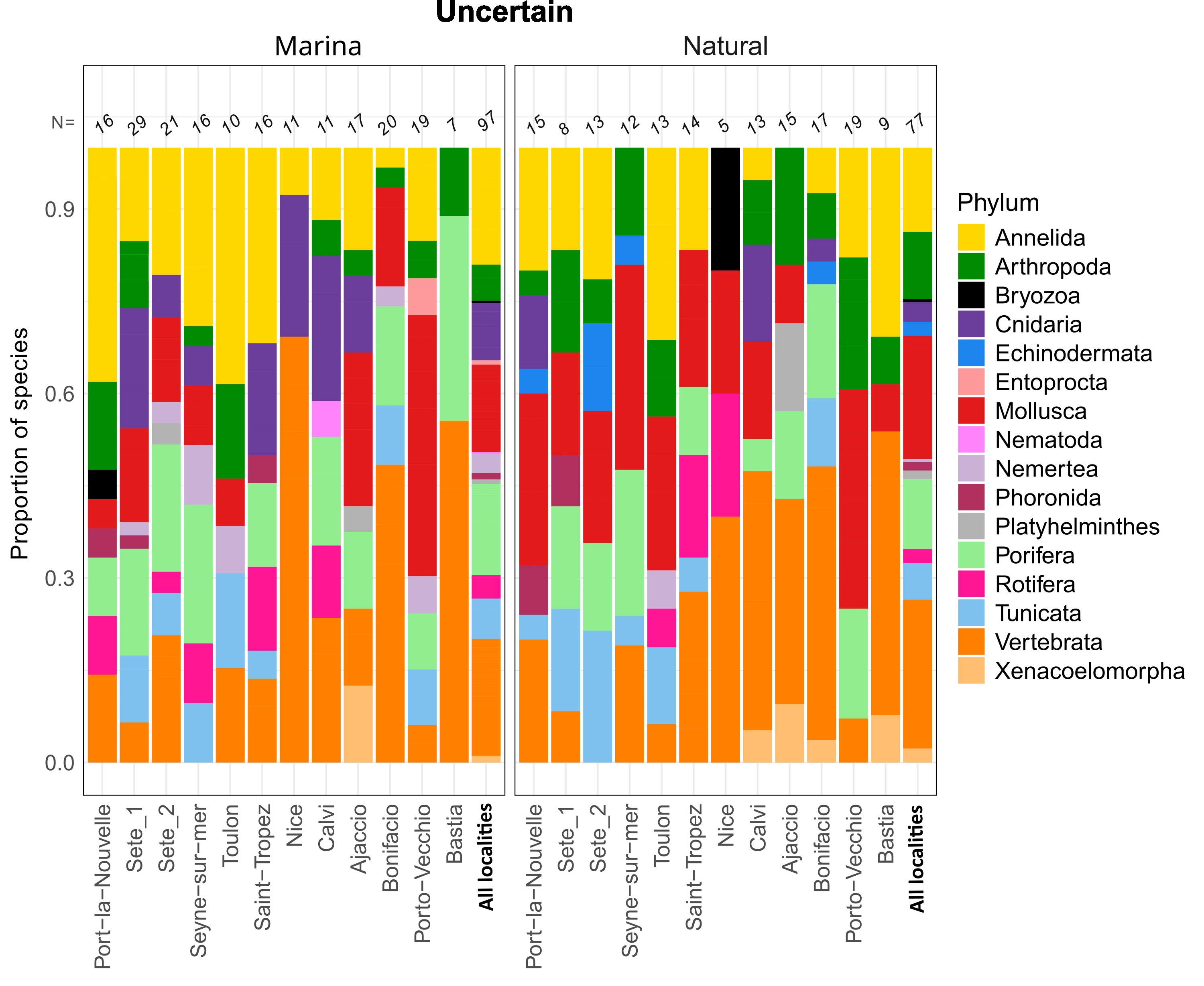

*Figure S2: Barplot displaying the proportion in the community of species included in the “Uncertain” category (i.e., species whose native range is not in the Mediterranean Sea or it is unknown, and that are not reported as NIS in Europe; see main text) by each habitat type and locality and across all localities. The number of Uncertain species recorded in the different habitat types is reported at the top of each bar. See Figure 1 in the main text for results for native species (NATS) and NIS.*

### Appendix S1.5: NIS proportion across localities

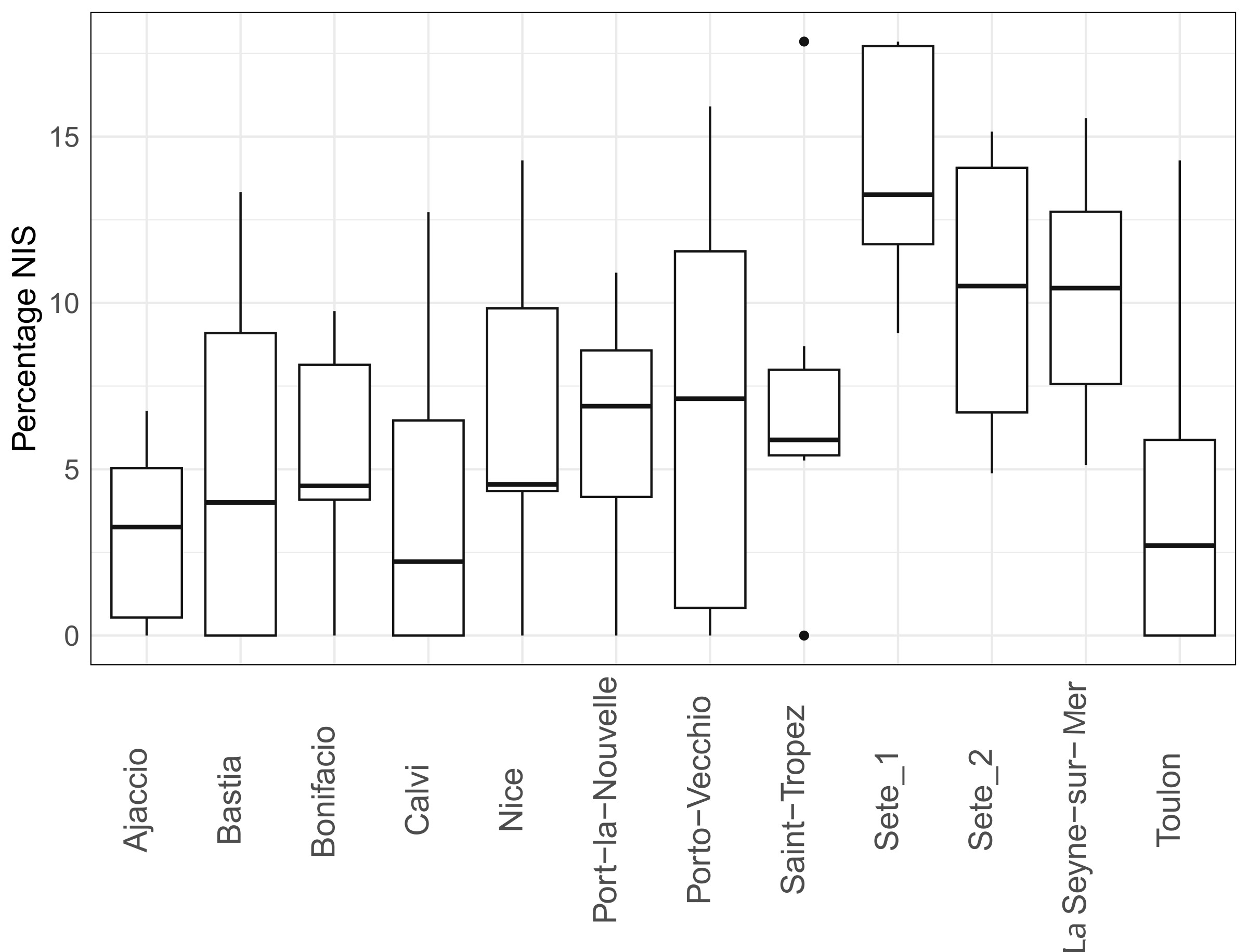
*Figure S3: Proportion of non-indigenous species (NIS) per locality combining both habitats (inside marinas and natural, i.e., 6 replicate samples per locality).*

### Appendix S1.6: Supplementary to beta diversity analyses

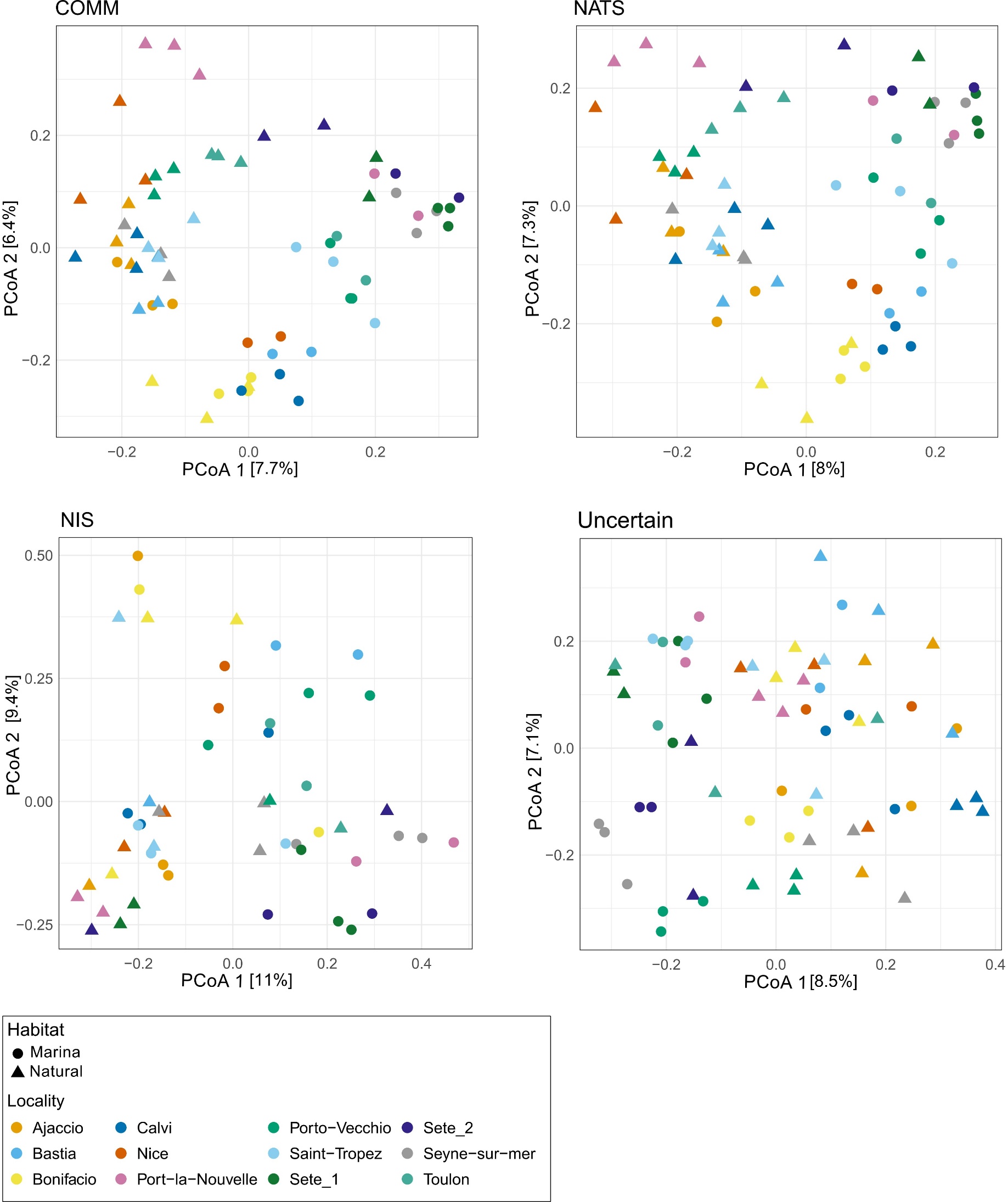

*Figure S4: Unconstrained ordination based on Principal Coordinate Analysis (PCoA) representing community dissimilarities calculated with Jaccard distance for the whole community (COMM, panel A), the native species (NATS, panel B), the Non-Indigenous Species (NIS, C) and species not included in any of the two former groups (Uncertain, D). Communities are coloured according to the locality in which they were sampled; the type of habitat is indicated by point shape.*

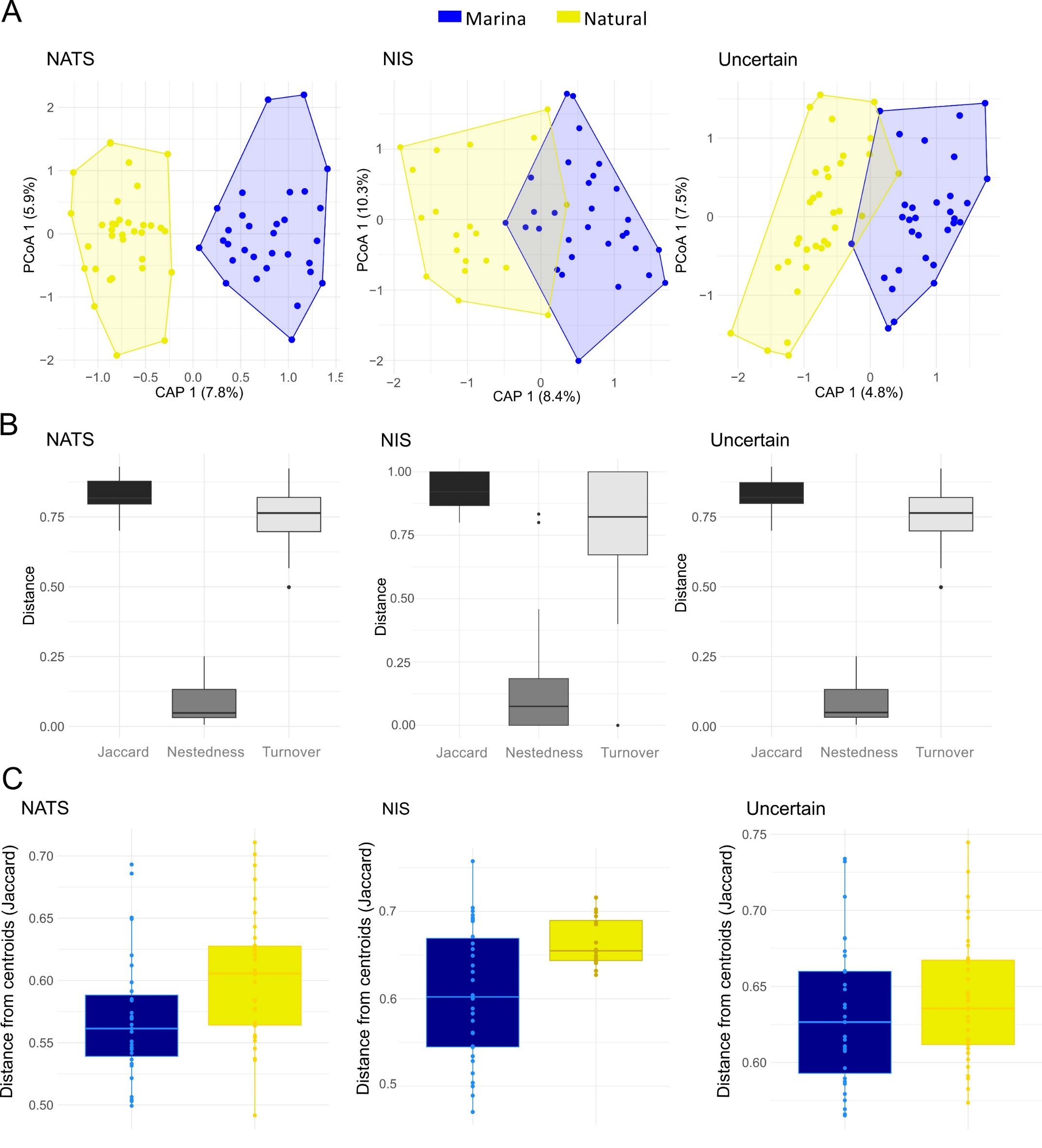
*Figure S5: A) Partial dbRDA performed to test the effect of habitat on NATS and NIS species composition, accounting for the effect of locality; B) boxplots displaying partitioning of Jaccard dissimilarities in its nestedness and turnover components with comparisons between samples from inside marinas and from natural sites; C)* *boxplot showing Jaccard dissimilarities between samples collected inside marinas (blue) and in natural habitats (yellow).*

### Appendix S1.7: NIS richness

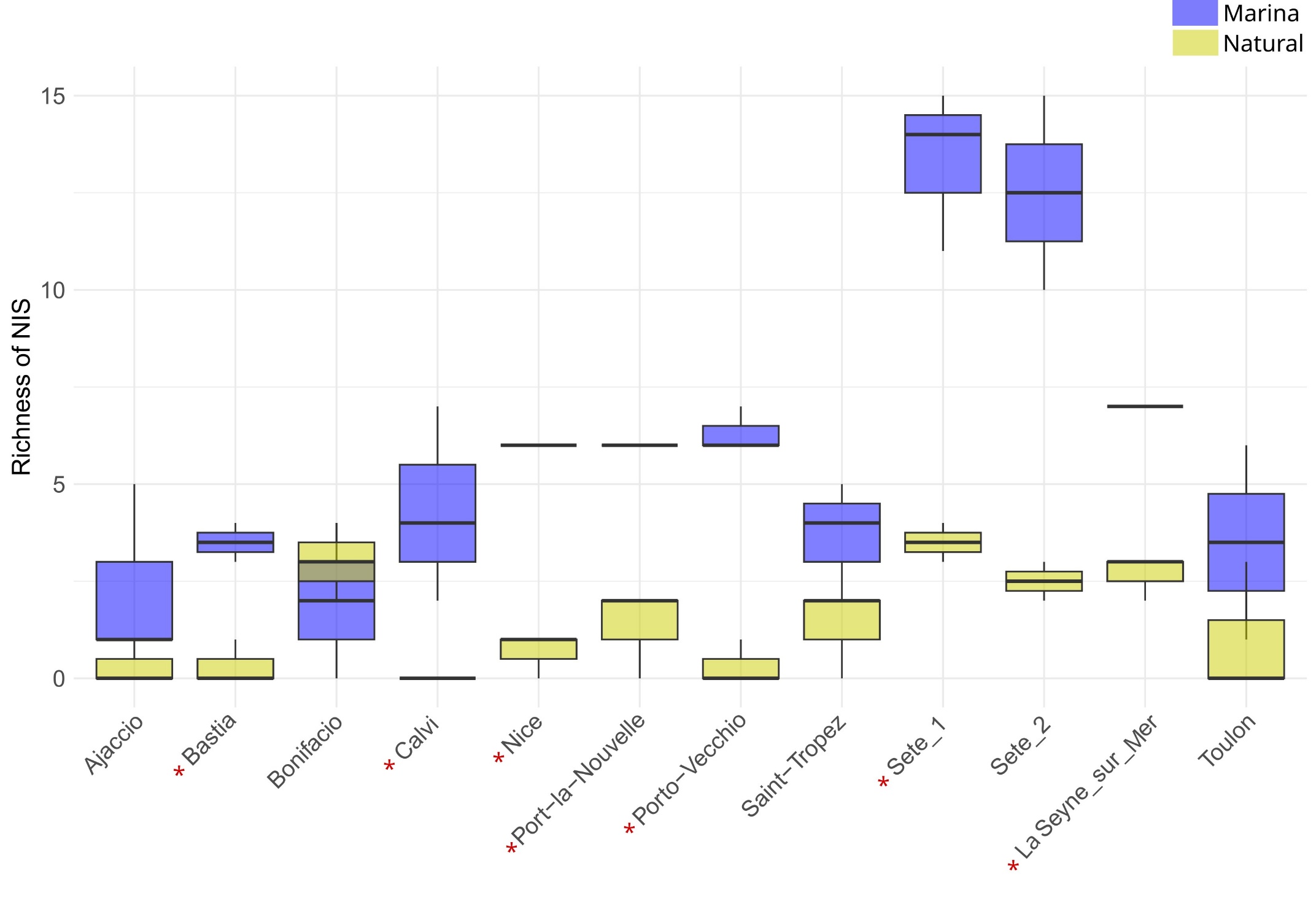

*Figure S6: Species richness of NIS communities inside marinas and in natural habitats across localities. Localities where the difference in NIS richness was significant are flagged respectively with red (Student’s t-test or Wilcoxon’s test, one-tailed, P value < 0.05).*

### Appendix S1.8: Species richness in marinas and in natural habitats

*Table S1:* *Generalized Linear Mixed Model's (GLMM) results describing the influence of habitat type on species richness of the whole (COMM), native (NATS), Non-Indigenous (NIS) and Uncertain communities. Standardised estimates are shown with their 95% confidence intervals, standard error, test statistics (Wald's) and p values. "Marina" was set as reference level used in the model.*

| **Category** | **Predictor** | **Estimate** | **95% CI** | **Std. Error** | **z value** | **p value** | **Ref.habitat** |
| --- | --- | --- | --- | --- | --- | --- | --- |
| COMM | Habitat type - Natural | -0.45 | [-0.58; -0.32] | 0.065 | -6.968 | < 0.00001 | Marina |
| NATS | Habitat type - Natural | -0.40 | [-0.54; -0.26] | 0.071 | -5.688 | < 0.00001 | Marina |
| NIS | Habitat type - Natural | -1.41 | [-1.74; -1.09] | 0.169 | -8.477 | < 0.00001 | Marina |
| Uncertain | Habitat type - Natural | -0.34 | [-0.51; -0.16] | 0.090 | -3.752 | 0.00018 | Marina |

*Table S2: Outputs of Student’s t-test (or Mann Whitney test, in case of data not normally distributed) performed to determine whether species richness (COMM) differed in marinas and in natural habitats in each locality. Significant results (P- value < 0.05) are reported in bold in the table.*

|  | **test** | **stat** | **p-value** |
| --- | --- | --- | --- |
| Ajaccio | t.test | 1.225 | 0.341 |
| Bastia | t.test | 4.666 | 0.091 |
| Bonifacio | wilcox | 3.500 | 0.825 |
| **Calvi** | **t.test** | **8.115** | **0.008** |
| Nice | wilcox | 6.000 | 0.149 |
| Port-la-Nouvelle | t.test | 2.767 | 0.094 |
| **La Seyne-sur-Mer** | **t.test** | **3.113** | **0.041** |
| Sete_1 | wilcox | 6.000 | 0.149 |
| Sete_2 | wilcox | 4.000 | 0.221 |
| Saint-Tropez | t.test | 0.055 | 0.959 |
| **Porto-Vecchio** | **t.test** | **3.843** | **0.019** |
| Toulon | t.test | 2.931 | 0.072 |

*Table S3: Generalized Linear Mixed Model's (GLMM) results describing the influence of habitat type on species richness in different phyla (COMM). Standardised estimates are shown alongside with 95% confidence intervals, standard error, test statistics (Wald's) and p values. "Marina" was set as reference level used in the model. GLMM with significant P-values (p value < 0.05) and with 95% confidence intervals not overlapping zero are shown in bold. Nematoda and Placozoa were excluded from this analysis due to not being recorded in more than one sample in the whole dataset.*

| **Predictor** | **Phylum** | **Estimate** | **95% CI** | **Std. Error** | **z value** | **Pr(>\|z\|)** | **Ref.habitat** |
| --- | --- | --- | --- | --- | --- | --- | --- |
| Habitat type - Natural | Echinodermata | 0.028 | [-0.484; 0.54] | 0.229 | 0.080 | 9.360E-01 | Marina |
|  | **Mollusca** | **-0.318** | **[-0.54; -0.096]** | **0.094** | **-3.310** | **9.334E-04** | **Marina** |
|  | **Arthropoda** | **0.356** | **[0.106; 0.606]** | **0.123** | **2.855** | **4.299E-03** | **Marina** |
|  | **Cnidaria** | **-0.814** | **[-1.22; -0.408]** | **0.188** | **-4.416** | **1.007E-05** | **Marina** |
|  | **Vertebrata** | **-0.537** | **[-0.706; -0.368]** | **0.062** | **-8.395** | **4.660E-17** | **Marina** |
|  | **Bryozoa** | **-1.962** | **[-2.904; -1.02]** | **0.481** | **-4.081** | **4.475E-05** | **Marina** |
|  | **Annelida** | **-0.721** | **[-1.019; -0.422]** | **0.119** | **-5.944** | **2.776E-09** | **Marina** |
|  | **Tunicata** | **-1.260** | **[-1.733; -0.786]** | **0.242** | **-5.213** | **1.854E-07** | **Marina** |
|  | Entoprocta | -17.870 | [-10487.61; 10451.869] | 1310.574 | -0.011 | 9.908E-01 | Marina |
|  | **Porifera** | **-0.509** | **[-0.856; -0.162]** | **0.176** | **-2.898** | **3.756E-03** | **Marina** |
|  | **Nemertea** | **-1.925** | **[-2.98; -0.869]** | **0.539** | **-3.575** | **3.508E-04** | **Marina** |
|  | Xenacoelomorpha | 0.431 | [-0.691; 1.553] | 0.572 | 0.753 | 4.514E-01 | Marina |
|  | Platyhelminthes | -0.467 | [-1.737; 0.802] | 0.648 | -0.722 | 4.704E-01 | Marina |
|  | Chaetognatha | 1.294 | [-0.897; 3.485] | 1.118 | 1.157 | 2.471E-01 | Marina |
|  | Phoronida | -0.010 | [-1.111; 1.09] | 0.561 | -0.019 | 9.852E-01 | Marina |
|  | **Rotifera** | **-1.115** | **[-2.139; -0.092]** | **0.522** | **-2.136** | **3.264E-02** | **Marina** |
|  | Nematoda | *NA* | *NA* | *NA* | *NA* | *NA* | *NA* |
|  | Placozoa | *NA* | *NA* | *NA* | *NA* | *NA* | *NA* |

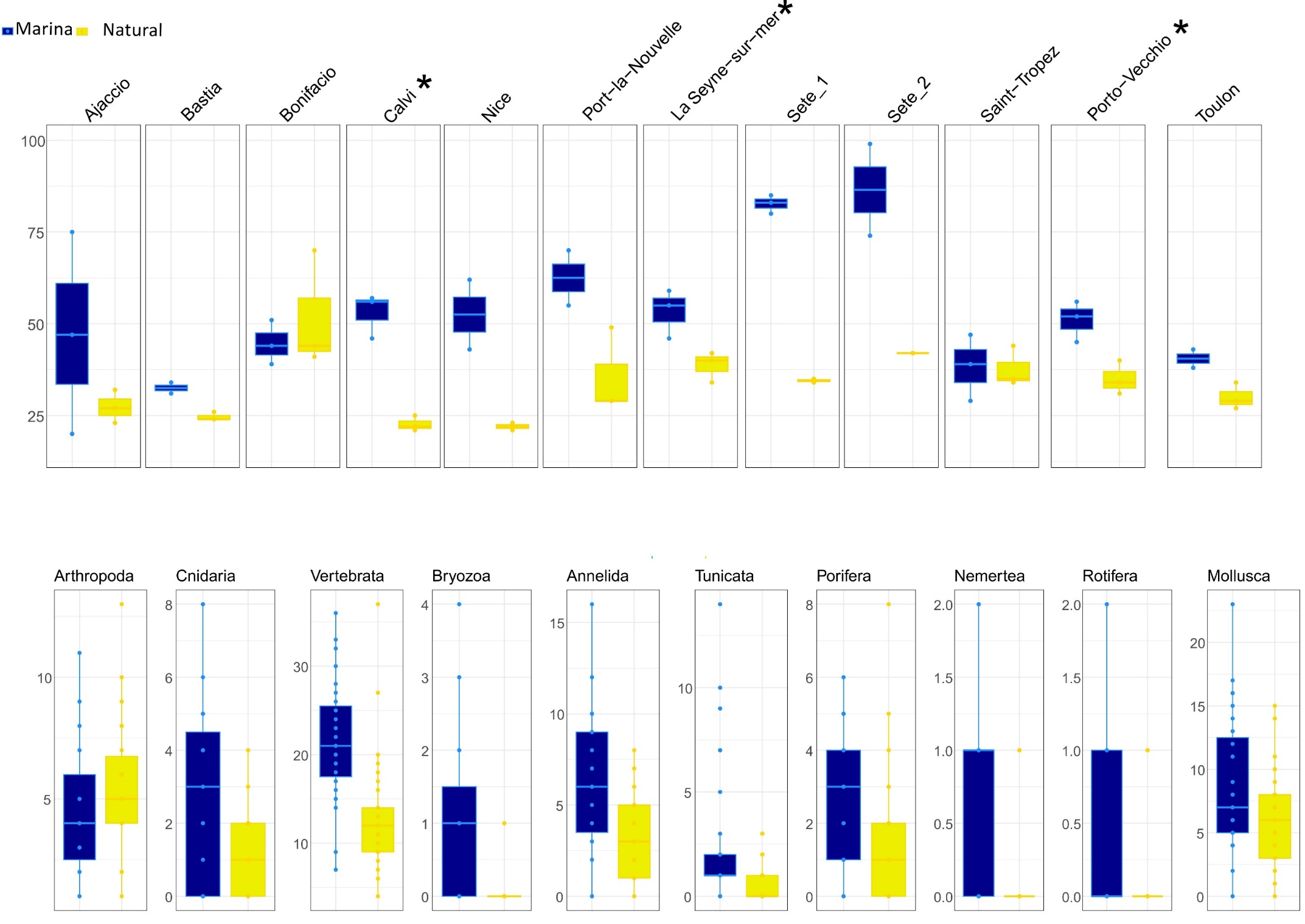

*Figure S7: Boxplots showing species richness (COMM) in marinas and natural habitats for each locality (top panel) and for each phylum (bottom panel). Localities showing significant difference of species richness between habitat types (Student’s t-test or Mann Whitney test, P value < 0.05) are flagged with an asterisk; in the bottom panel, only phyla whose species richness differed significantly between the two habitat types according to GLMMs are reported.*

### Appendix S1.9: Indicator species, and their taxonomic and functional traits, behind communities dissimilarities in marinas and in natural habitats

*Table S4: Species significantly associated with marinas or natural habitats according to Indicator Value Analysis* *performed by indicspecies package in R (Indval.g, P value < 0.05). NIS indicator species are reported in red.*

|  | **p.value** | **stat** | **Kingdom** | **Phylum** | **Class** | **Order** | **Family** | **Genus** | **Species** |
| --- | --- | --- | --- | --- | --- | --- | --- | --- | --- |
| **Natural** | 0.001 | 0.715 | Animalia | Arthropoda | Copepoda | Calanoida | Paracalanidae | Paracalanus | *Paracalanus quasimodo* |
|  | 0.004 | 0.652 | Animalia | Arthropoda | Copepoda | Calanoida | Acartiidae | Acartia | *Acartia (Acartiura) clausii* |
|  | 0.001 | 0.647 | Animalia | Arthropoda | Copepoda | Calanoida | Centropagidae | Centropages | *Centropages typicus* |
|  | 0.047 | 0.461 | Animalia | Vertebrata | Teleostei | Clupeiformes | Dorosomatidae | Sardinella | *Sardinella longiceps* |
|  | 0.029 | 0.455 | Animalia | Vertebrata | Teleostei | Beloniformes | Belonidae | Belone | *Belone belone* |
|  | 0.045 | 0.422 | Animalia | Vertebrata | Teleostei | Eupercaria incertae sedis | Labridae | Symphodus | *Symphodus ocellatus* |
|  | 0.038 | 0.42 | Animalia | Mollusca | Bivalvia | Cardiida | Tellinidae | Scissula | *Scissula similis* |
| **Marina** | 0.001 | 0.739 | Animalia | Vertebrata | Teleostei | Mugiliformes | Mugilidae | Chelon | *Chelon ramada* |
|  | 0.001 | 0.726 | Animalia | Annelida | Polychaeta | Sabellida | Serpulidae | Vermiliopsis | *Vermiliopsis striaticeps* |
|  | 0.001 | 0.699 | Animalia | Porifera | Demospongiae | Haplosclerida | Callyspongiidae | Callyspongia | *Callyspongia (Cladochalina) plicifera* |
|  | 0.003 | 0.698 | Animalia | Vertebrata | Teleostei | Mulliformes | Mullidae | Mullus | *Mullus surmuletus* |
|  | 0.001 | 0.677 | Animalia | Vertebrata | Teleostei | Mugiliformes | Mugilidae | Chelon | *Chelon auratus* |
|  | 0.001 | 0.672 | Animalia | Vertebrata | Teleostei | Gobiiformes | Gobiidae | Gobius | *Gobius cobitis* |
|  | 0.004 | 0.624 | Animalia | Vertebrata | Teleostei | Eupercaria incertae sedis | Sparidae | Pagellus | *Pagellus erythrinus* |
|  | 0.001 | 0.622 | Animalia | Tunicata | Ascidiacea | Stolidobranchia | Styelidae | Botryllus | *Botryllus schlosseri* |
|  | 0.006 | 0.607 | Animalia | Vertebrata | Teleostei | Mulliformes | Mullidae | Mullus | *Mullus barbatus* |
|  | 0.002 | 0.606 | Animalia | Nemertea | Palaeonemertea | Archinemertea | Cephalotrichidae | Cephalothrix | *Cephalothrix rufifrons* |
|  | 0.002 | 0.606 | Animalia | Cnidaria | Hydrozoa | Anthoathecata | Eudendriidae | Eudendrium | *Eudendrium racemosum* |
|  | 0.002 | 0.606 | Animalia | Mollusca | Bivalvia | Ostreida | Ostreidae | Ostrea | *Ostrea stentina* |
|  | 0.002 | 0.58 | Animalia | Annelida | Polychaeta | Terebellida | Terebellidae | Terebella | *Terebella lapidaria* |
|  | 0.001 | 0.572 | Animalia | Mollusca | Bivalvia | Ostreida | Ostreidae | Ostrea | *Ostrea edulis* |

*continues*

|  | **p.value** | **stat** | **Kingdom** | **Phylum** | **Class** | **Order** | **Family** | **Genus** | **Species** |
| --- | --- | --- | --- | --- | --- | --- | --- | --- | --- |
| **Marina** | 0.001 | 0.568 | Animalia | Tunicata | Ascidiacea | Stolidobranchia | Pyuridae | Microcosmus | *Microcosmus squamiger* |
|  | 0.007 | 0.561 | Animalia | Mollusca | Bivalvia | Ostreida | Ostreidae | Saccostrea | *Saccostrea glomerata* |
|  | 0.004 | 0.544 | Animalia | Annelida | Polychaeta | Spionida | Spionidae | Pseudopolydora | *Pseudopolydora paucibranchiata* |
|  | 0.001 | 0.539 | Animalia | Bryozoa | Gymnolaemata | Cheilostomatida | Cryptosulidae | Cryptosula | *Cryptosula pallasiana* |
|  | 0.008 | 0.522 | Animalia | Annelida | Polychaeta | Phyllodocida | Hesionidae | Syllidia | *Syllidia armata* |
|  | 0.042 | 0.516 | Animalia | Arthropoda | Copepoda | Calanoida | Pseudodiaptomidae | Pseudodiaptomus | *Pseudodiaptomus marinus* |
|  | 0.022 | 0.503 | Animalia | Porifera | Demospongiae | Suberitida | Halichondriidae | Halichondria | *Halichondria (Halichondria) bowerbanki* |
|  | 0.023 | 0.491 | Animalia | Vertebrata | Teleostei | Gadiformes | Phycidae | Phycis | *Phycis phycis* |
|  | 0.007 | 0.481 | Animalia | Mollusca | Gastropoda | NA | Limapontiidae | Limapontia | *Limapontia capitata* |
|  | 0.008 | 0.481 | Animalia | Vertebrata | Teleostei | Gobiiformes | Gobiidae | Pomatoschistus | *Pomatoschistus minutus* |
|  | 0.002 | 0.475 | Animalia | Tunicata | Ascidiacea | Stolidobranchia | Styelidae | Styela | *Styela plicata* |
|  | 0.003 | 0.475 | Animalia | Mollusca | Bivalvia | Cardiida | Semelidae | Abra | *Abra alba* |
|  | 0.003 | 0.475 | Animalia | Annelida | Polychaeta | Sabellida | Sabellidae | Parasabella | *Parasabella saxicola* |
|  | 0.006 | 0.475 | Animalia | Nemertea | Palaeonemertea | Archinemertea | Cephalotrichidae | Cephalothrix | *Cephalothrix hongkongiensis* |
|  | 0.036 | 0.472 | Animalia | Vertebrata | Teleostei | Mugiliformes | Mugilidae | Chelon | *Chelon saliens* |
|  | 0.033 | 0.458 | Animalia | Vertebrata | Teleostei | Gobiiformes | Gobiidae | Zosterisessor | *Zosterisessor ophiocephalus* |
|  | 0.038 | 0.458 | Animalia | Vertebrata | Teleostei | Eupercaria incertae sedis | Sparidae | Oblada | *Oblada melanurus* |
|  | 0.021 | 0.447 | Animalia | Vertebrata | Teleostei | Eupercaria incertae sedis | Sparidae | Diplodus | *Diplodus annularis* |
|  | 0.027 | 0.447 | Animalia | Cnidaria | Hydrozoa | Leptothecata | Kirchenpaueriidae | Kirchenpaueria | *Kirchenpaueria halecioides* |
|  | 0.033 | 0.447 | Animalia | Vertebrata | Teleostei | Gobiiformes | Gobiidae | Gobius | *Gobius cruentatus* |

*continues*

|  | **p.value** | **stat** | **Kingdom** | **Phylum** | **Class** | **Order** | **Family** | **Genus** | **Species** |
| --- | --- | --- | --- | --- | --- | --- | --- | --- | --- |
| **Marina** | 0.008 | 0.44 | Animalia | Cnidaria | Hydrozoa | Anthoathecata | Tubulariidae | Ectopleura | *Ectopleura marina* |
|  | 0.01 | 0.44 | Animalia | Vertebrata | Teleostei | Gadiformes | Merlucciidae | Merluccius | *Merluccius merluccius* |
|  | 0.046 | 0.41 | Animalia | Annelida | Polychaeta | Sabellida | Serpulidae | Serpula | *Serpula columbiana* |
|  | 0.047 | 0.41 | Animalia | Vertebrata | Teleostei | Eupercaria incertae sedis | Sparidae | Diplodus | *Diplodus sargus* |
|  | 0.019 | 0.402 | Animalia | Mollusca | Polyplacophora | Chitonida | Tonicellidae | Lepidochitona | *Lepidochitona cinerea* |
|  | 0.02 | 0.402 | Animalia | Porifera | Demospongiae | Suberitida | Suberitidae | Protosuberites | *Protosuberites mereui* |
|  | 0.021 | 0.402 | Animalia | Annelida | Polychaeta | Spionida | Spionidae | Malacoceros | *Malacoceros fuliginosus* |
|  | 0.021 | 0.402 | Animalia | Bryozoa | Gymnolaemata | Cheilostomatida | Watersiporidae | Watersipora | *Watersipora subtorquata* |
|  | 0.023 | 0.402 | Animalia | Cnidaria | Hexacorallia | Actiniaria | Edwardsiidae | Edwardsia | *Edwardsia longicornis* |
|  | 0.025 | 0.402 | Animalia | Cnidaria | Scyphozoa | Semaeostomeae | Ulmaridae | Aurelia | *Aurelia coerulea* |
|  | 0.027 | 0.402 | Animalia | Vertebrata | Teleostei | Mugiliformes | Mugilidae | Mugil | *Mugil cephalus* |
|  | 0.029 | 0.402 | Animalia | Annelida | Polychaeta | Echiuroidea | Bonelliidae | Bonellia | *Bonellia viridis* |
|  | 0.042 | 0.359 | Animalia | Annelida | Polychaeta | Sabellida | Serpulidae | Hydroides | *Hydroides elegans* |
|  | 0.043 | 0.359 | Animalia | Tunicata | Ascidiacea | Stolidobranchia | Styelidae | Polycarpa | *Polycarpa pomaria* |
|  | 0.044 | 0.359 | Animalia | Annelida | Polychaeta | NA | Paraonidae | Paradoneis | *Paradoneis ilvana* |
|  | 0.044 | 0.359 | Animalia | Vertebrata | Teleostei | Clupeiformes | Clupeidae | Sprattus | *Sprattus sprattus* |
|  | 0.045 | 0.359 | Animalia | Bryozoa | Stenolaemata | Cyclostomatida | Crisiidae | Filicrisia | *Filicrisia geniculata* |
|  | 0.047 | 0.359 | Animalia | Vertebrata | Teleostei | Lophiiformes | Lophiidae | Lophius | *Lophius budegassa* |
|  | 0.05 | 0.359 | Animalia | Arthropoda | Malacostraca | Decapoda | Carcinidae | Carcinus | *Carcinus maenas* |

*Table S5: Outputs of tests performed to assess the relationship between species (COMM, NATS, NIS and Uncertain) scores distribution on the first axis of the partial dbRDA and their taxonomical and functional traits. For COMM, the classification as NIS, NATS or Uncertain was also used to explain species scores distribution (i.e. origin category). For NIS, ecology traits could not be tested on fish due to the limited number of species recorded for this group (N=2).*

| **Group** | **Trait** | **test** | **stat** | **p value** | **Significant Dunn Post-hoc test (p value "bh" adjusted < 0.05)** |
| --- | --- | --- | --- | --- | --- |
| COMM | Motility | Wilcoxon-Mann-Whitney | 26490 | **0.002** |  |
|  | Ecology -Non fishes |  | 11452.5 | **3.1E-06** |  |
|  | Ecology - Fishes | Kruskal-Wallis | 8.3 | **0.039** | *reef associated ≠ bethopelagic (p=0.04), reef associated ≠ demersal (p=0.05)* |
|  | Phylum |  | 42.6 | **0.0005** | *Arthropoda* ≠ *Tunicata (p=0.003),Echinodermata ≠ Tunicata (p=0.01), Arthropoda ≠ Vertebrata (p=0.005)* |
|  | Class |  | 57.4 | **0.016** | *Ascidiacea ≠ Copepoda (p=0.005), Copepoda ≠ Teleostei (p=0.01)* |
|  | Fishing vulnerability | Spearman’s_correlation | 478379.6 | 0.067 |  |
|  | Category (NIS, NATS, Uncertain) | Kruskal-Wallis | 12.1 | **0.002** | *NIS ≠ NATS (p= 0.0008), NIS ≠ Uncertain (p=0.0013)* |
| NATS | Motility | Wilcoxon-Mann-Whitney | 10217.5 | 0.073 |  |
|  | Ecology -Non fishes |  | 5210.5 | **2.3E-07** |  |
|  | Ecology - Fishes | Kruskal-Wallis | 5.3 | 0.150 |  |
|  | Phylum |  | 38.6 | **0.001** | *Arthropoda ≠ Vertebrata (p=0.0003)* |
|  | Class |  | 57.0 | **0.011** | *Copepoda ≠ Teleostei (p=0.0002), Ascidiacea ≠ Copepoda (p=0.05)* |
|  | Fishing vulnerability | Spearman’s_correlation | 234291.7 | 0.189 |  |
| NIS | Motility | Wilcoxon-Mann-Whitney | 180.5 | 0.1879 |  |
|  | Ecology -Non fishes |  | 157.0 | 0.9448 |  |
|  | Ecology - Fishes* | Kruskal-Wallis | *NA* | *NA* |  |
|  | Phylum |  | 14.4 | 0.043 | *No significant pairwise comparisons* |
|  | Class |  | 20.7 | 0.078 |  |
|  | Fishing vulnerability | Not possible to perform test, only 2 fishes | *NA* | *NA* |  |
| Uncertain | Motility | Wilcoxon-Mann-Whitney | 1786.5 | 0.313 |  |
|  | Ecology -Non fishes |  | 479.5 | 0.107 |  |
|  | Ecology - Fishes | Kruskal-Wallis | 2.2 | 0.542 |  |
|  | Phylum |  | 17.1 | 0.313 |  |
|  | Class |  | 25.4 | 0.230 |  |
|  | Fishing vulnerability | Spearman`s_correlation | 3843.8 | 0.224 |  |
